# Hemispheric asymmetry of globus pallidus relates to alpha modulation in reward-related attentional tasks

**DOI:** 10.1101/303016

**Authors:** C. Mazzetti, T. Staudigl, T. R. Marshall, J. M. Zumer, S. J. Fallon, O. Jensen

**Affiliations:** Radboud University Nijmegen, Donders Institute for Brain, Cognition and Behaviour, Nijmegen 6525EN, The Netherlands; Ludwig-Maximilians University of Munich, Department of Psychology, 80539, Munich, Germany; Computational Cognitive Neuroscience Lab, Department of Experimental Psychology, University of Oxford, OX2 6GG, Oxford, UK; School of Life and Health Sciences, Aston University, B4 7ET, Birmingham, UK; National Institute for Health Research, Bristol Biomedical Research Centre, University Hospitals Bristol NHS Foundation Trust and University of Bristol, BS1 3NU, Bristol, UK; Centre for Human Brain Health, School of Psychology, University of Birmingham, B15 2TT Birmingham, UK

## Abstract

While subcortical structures like the basal ganglia have been widely explored in relation to motor control, recent evidence suggests that their mechanisms extend to the domain of attentional switching. We here investigated the subcortical involvement in reward related top-down control of visual alpha-band oscillations (8 – 13 Hz), which have been consistently linked to mechanisms supporting the allocation of visuo-spatial attention. Given that items associated with contextual saliency (e.g. monetary reward or loss) attract attention, it is not surprising that the acquired salience of visual items further modulates. The executive networks controlling such reward-dependent modulations of oscillatory brain activity have yet to be fully elucidated. Although such networks have been explored in terms of cortico-cortical interactions, subcortical regions are likely to be involved. To uncover this, we combined MRI and MEG data from 17 male and 11 female participants, investigating whether derived measures of subcortical structural asymmetries predict interhemispheric modulation of alpha power during a spatial attention task. We show that volumetric hemispheric lateralization of globus pallidus (GP) and thalamus (Th) explains individual hemispheric biases in the ability to modulate posterior alpha power. Importantly, for the GP, this effect became stronger when the value-saliency parings in the task increased. Our findings suggest that the GP and Th in humans are part of a subcortical executive control network, differentially involved in modulating posterior alpha activity in the presence of saliency. Further investigation aimed at uncovering the interaction between subcortical and neocortical attentional networks would provide useful insight in future studies.

**Significance statement:** While the involvement of subcortical regions into higher level cognitive processing, such as attention and reward attribution, has been already indicated in previous studies, little is known about its relationship with the functional oscillatory underpinnings of said processes. In particular, interhemispheric modulation of alpha band (8-13Hz) oscillations, as recorded with magnetoencephalography (MEG), has been previously shown to vary as a function of salience (i.e. monetary reward/loss) in a spatial attention task. We here provide novel insights into the link between subcortical and cortical control of visual attention. Using the same reward-related spatial attention paradigm, we show that the volumetric lateralization of subcortical structures (specifically Globus Pallidus and Thalamus) explains individual biases in the modulation of visual alpha activity.

## Introduction

Functioning in the natural world necessitates the presence of neuronal mechanisms capable of prioritising stimuli according to their relevance (Nobre and Kastner, 2014). Deployment of attentional resources is biased towards stimuli associated with salience (e.g. monetary reward or loss), even when unrelated to the current task (Chelazzi et al., 2013). Posterior neuronal oscillations in the alpha band (8 – 13 Hz) reflect the allocation of covert attention (Worden et al., 2000; Kelly, 2006; Thut, 2006; Jensen and Mazaheri, 2010) and they have been shown to be mediated by cortico-cortical interactions (Capotosto et al., 2012a; Ptak, 2012; Vossel et al., 2014; Marshall et al., 2015a, 2015b). On the other hand, these cortical networks are further modulated by subcortical input (van Schouwenburg et al., 2010a, 2010b), whose involvement in posterior oscillations remains still unclear. Previous literature has indeed linked subcortical activity to cognitive control (Cummings, 1993; Jahfari et al., 2011; Braunlich and Seger, 2013), but a direct link between these structures and alpha band oscillations has not been established.

Electrophysiological activity from subcortical regions are poorly detected with magnetoencephalography (MEG). Alternatively, subcortical structures measured by magnetic resonance imaging (MRI) can be related to oscillatory brain activity (Tomer et al., 2008, 2013). For instance, it has been demonstrated that individual hemispheric asymmetries in the volume of the superior longitudinal fasciculus (SLF) relates to the individual ability to modulate posterior alpha oscillations (Marshall et al., 2015a). Importantly, subjects with greater right than left SLF volume also displayed higher modulation of posterior alpha activity in the left hemisphere, compared to the right (and vice versa). Through an analogous approach, we postulated that volumetric asymmetries of subcortical areas would be reflected by individual interhemispheric biases in the modulation of alpha oscillations during selective attention in a reward context. Basal ganglia (BG), in addition to motor control, have a well-established role in reward processing and salience attribution (Hikosaka et al., 2008, 2014; Shulman et al., 2010a; Braunlich and Seger, 2013), and recent studies have already pointed to their functions extending into higher level cognitive processing (Arcizet and Krauzlis, 2018). This notion has been initially explored in animal recordings (Tremblay et al., 1998; Schultz et al., 2000; Lauwereyns et al., 2002; Shipp, 2004; Saalmann and Kastner, 2011; Schechtman et al., 2016), while in humans, it has recently been suggested that the BG play also a specific role in spatial attention and selection (van Schouwenburg et al., 2010a; Tommasi et al., 2014; Van Schouwenburg et al., 2015). Another subcortical structure playing a crucial role in cognitive processing is the thalamus (Fiebelkorn et al., 2019; Jaramillo et al., 2019), whose nuclei are involved in the regulation of synchronized activity in the visual cortex in relation to visual attention and largely interact with the BG (Lopes da Silva et al., 1980; Saalmann et al., 2012; Zhou et al., 2016; Halgren et al., 2017).

We here re-analysed MEG and structural data collected in a previous study which considered the impact of stimuli paired with value-salience on the modulation of oscillatory brain activity in a covert attention task (Marshall et al., 2017). The participants performed a spatial cueing task, with Chinese symbols serving as targets and distractors. Prior to the recordings, stimuli were paired with monetary rewards or losses. Marshall et al. successfully demonstrated a location-specific influence for the stimuli associated with reward and loss. Specifically, alpha lateralization demonstrated sensitivity to stimulus salience, but not to stimulus valence: both positive and negative targets (i.e. salient targets) produced increased alpha lateralisation compared to neutral targets, and both positive and negative distractors (i.e. salient distractors) reduced alpha lateralisation compared to neutral distractors. Given these findings, we here examine the further link between lateralization of subcortical structures and alpha oscillations. We hence re-analysed these data with the aim to investigate the putative role of the subcortical brain areas in biasing alpha power modulation during attentional shifts to stimuli paired with value-saliency. MRI data of the participants were processed in order to estimate volumetric asymmetries of subcortical areas, consistent with methods employed in previous studies on clinical and healthy population (Womer et al., 2014; Guadalupe et al., 2016; Okada et al., 2016). We focused on identifying the link between individual volumetric asymmetries of subcortical areas and individual interhemispheric bias in the ability to modulate posterior alpha oscillations. Crucially, we further examined whether this relationship was affected by the degree of stimulus-value associations in the task. Furthermore, we included 612 MRI scans to evaluate subcortical volumetric asymmetries in a larger dataset.

## Materials and Methods

### Participants

In the present study, we re-analysed the previously acquired dataset described in (Marshall et al., 2017), where twenty-eight healthy volunteers participated in the study (mean age: 23±2.7 years; 17 female; all right handed). All participants reported normal or corrected-to-normal vision and no prior knowledge of Chinese language. Of these, datasets from three participants were excluded from the analysis (due to respectively: technical error during acquisition, excessive eye movements during MEG recording, and structural MRI data not acquired), leaving 25 participants. The experiment was conducted in compliance with the Declaration of Helsinki and was approved by the local ethics board (CMO region Arnhem-Nijmegen, CMO2001/095).

### Experimental design

The experiment consisted of two phases: in the learning phase, participants were trained to memorize associations between 6 Chinese characters and 3 different values (positive, negative, neutral). Conditioning was implemented by means of visual and auditory feedback: two symbols were associated with reward (+80 cents and a ‘kaching’ sound), two with loss (−80 cents and a ‘buzz’ sound) and two with no value (0 cents and a ‘beep’ sound) (see Figure 1A for an example stimulus-reward association). The stimulus-reward pairing was randomized across participants. Each trial started with the display of three fixation crosses (1000ms), followed by the presentation of a Chinese character (1000ms), together with its matching visual and auditory feedback (Figure 1B). Stimuli were displayed on a grey background, each of them was presented twelve times in a randomized order. The learning phase was conducted in a laboratory with attenuated sound and light and without MEG recording. With the aim of reducing extinction, upon completion of this phase participants were informed that the learnt stimulus-feedback associations would be signalling real reward outcomes throughout the testing phase (i.e. the presentation of a Chinese character, irrespective of its role as target or distractor, would result in a financial reward, loss or none). After the learning phase, participants performed a testing phase (Figure 1C), when they were required to perform a covert spatial attention tasks including the stimuli previously associated with a monetary outcome, while ongoing electromagnetic activity was recorded with MEG. In the testing phase, participants performed 8 blocks of 72 trials. Each trial started with the presentation of three fixation crosses for 1000 ms (pre-trial interval), whose contrast subsequently decreased, as a preparatory cue indicating imminent stimuli presentation. After 500ms, two symbols were presented to the left and right of the screen (8 degrees visual angle) respectively, together with a central fixation cross flanked by two arrows, indicating the target side. Participants were instructed to covertly attend the symbol on the cued side (‘target’) and to ignore the other one (‘distractor’), until one of them changed contrast. The contrast change either increased or decreased with equal probability, with onset after 750 ms (13% trials), 1450 ms (47% trials, ‘short interval trials’) or 2350 ms (40%, ‘long interval trials’) from stimulus presentation. Participants were asked to report the direction of the contrast change at the targeted (‘cued’) location as quickly as possible by button press, using the index or middle finger of the right hand to indicate their choice (finger-direction mapping was randomized across participants). Participants were instructed to refrain from responding when the distractor changed contrast. Shorter intervals of 750 ms were used to ensure that participants would start covertly directing their attention rapidly after the cue; these trials were not included in the analysis. The target changed contrast on 95% of the trials (valid trials), whereas in the remaining trials the distractor did (invalid trials). The approximate duration of the full task in the MEG was 50 minutes.

**Figure 1.**
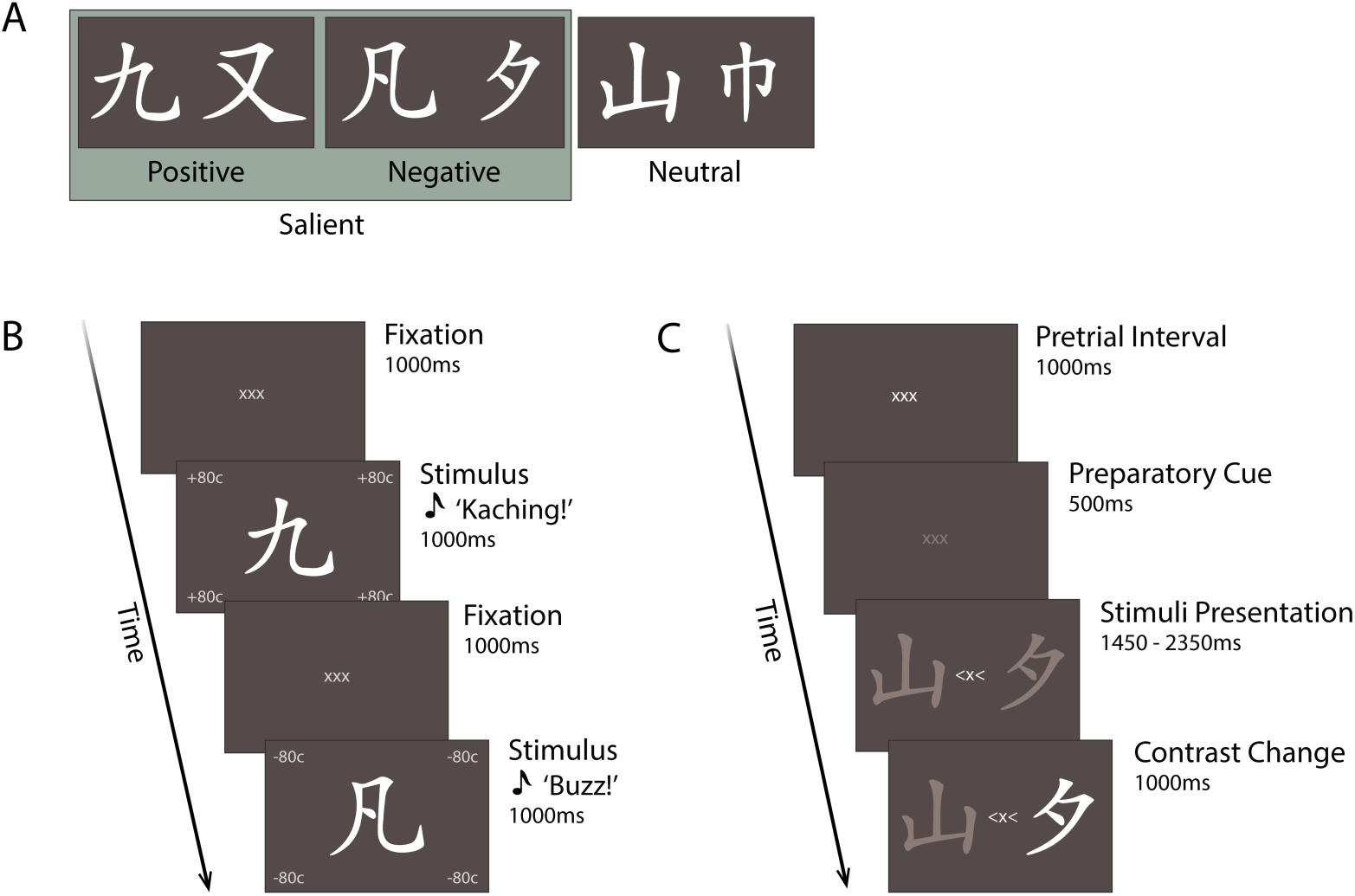
Illustration of selective attention task: stimuli and reward manipulation. (A) Six Chinese symbols served as stimuli for the task and were associated with three values: two paired with reward, two with loss and two with no financial change (neutral). (B) Representative trial of the learning phase. Symbols were displayed for 1000ms, systematically paired with the corresponding (positive, negative or neutral) value, via visual and auditory feedback. Characters presentation was alternated with a 1000ms fixation period. During the training phase, participants learned associations between the stimuli and their reward value. **C.** Representative trial of the testing phase. After a 1000ms pretrial interval, participants were primed with a 500ms preparatory cue signalling the upcoming stimuli. Two characters were then presented to the left and right hemifield, together with a spatial cue, instructing participants to covertly attend the symbol on the cued side (target) and ignore the other one (distractor). Participants’ task was to report when the target stimulus changed contrast. Contrast change could either occur after 750ms (13% of trials), 1450ms (47% of trials) or 2350ms (40% of the trials). In 95% of the trials, the target changed contrast (valid trials), whilst in 5% of the trials, the distractor changed contrast (invalid trials). Figure adapted from (Marshall et al., 2017).

As a result of the conditioning manipulation in the learning phase, targets and distractors in the task would be associated with either a salient (positive or negative) or a neutral value, resulting in three categories of trials of interest, as represented by different levels of value-salience, namely: zero (target and distractor neutral), one (target or distractor salient) or two (target and distractor salient) value-salience levels.

### MEG data acquisition

Electromagnetic brain activity was recorded from participants while seated, using a CTF 275-channels whole-head MEG system with axial gradiometers (CTF MEG Systems, VSM MedTech Ltd.). The data were sampled at 1200Hz, following an antialiasing filter set at 300Hz. Head position was constantly monitored throughout the experiment via online head-localization software. This had access to the position of the three head localization coils placed at anatomical fiducials (nasion, left and right ear), allowing, if necessary, readjustment of the participant’s position between blocks (Stolk et al., 2013). Horizontal and vertical EOG and ECG electrodes were recorded with bipolar Ag/AgCl electrodes.

### MEG data analysis

MEG data analysis was performed using the *FieldTrip* Toolbox running in MATLAB (Oostenveld et al., 2011). Continuous data were segmented in epochs, centred at the onset of the target contrast change, encompassing the preceding 1500 ms and the following 200 ms (this way covering the full stimulus presentation window for short trials). A notch filter was applied at 50, 100, 150 Hz to remove line noise, the mean was subtracted and the linear trend removed. Automatic artifact rejection was implemented for detection and removal of trials containing eye blinks and horizontal eye movements (detected with EOG), MEG sensor jumps and muscle artifacts. We produced virtual planar gradiometers by computing spatial derivatives of the magnetic signal recorded with axial gradiometers (Bastiaansen and Knösche, 2000). The method has the advantage of improving the interpretation of the topographic mapping since neural sources would produce a gradient field directly above them. Time-frequency representations (TFR) of power were then calculated for the resulting pairs of orthogonal planar gradiometers, before summing the power values at each sensor.

The analysis was performed by sliding a fixed time window of 500 ms in steps of 50 ms. The resulting data segments were multiplied by a Hanning taper and a fast Fourier transform was applied in the 2 – 30Hz frequency range, in steps of 2Hz. This procedure was applied only for correct valid trials, separately for left and right cued conditions.

For each participant, TFRs were averaged across trials and a Modulation Index (MI) was computed for each sensor *k* and over all time points *t* belonging to the time window of interest −750 – 0 ms, according to the formula:

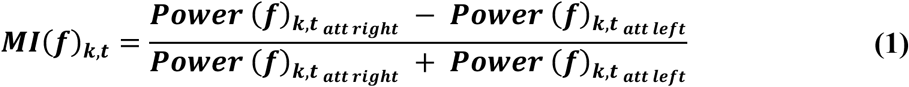

Where *Power*(*f*)*_k,t_att left__* represents the power at a given frequency *f* in the condition ‘attend left’ and *Power*(*f*)*_k,t_att right__* is the power of the same frequency in the condition ‘attend right’. As a result, positive (or negative) MI values, at a given sensor *k* and given timepoint *t*, indicate higher power at a given frequency *f* when attention was covertly directed towards the right (or left) hemifield.

Two clusters of sensors were then derived, by selecting the twenty symmetrical occipito-parietal sensors (i.e. ten pairs of sensors) showing the highest interhemispheric difference in alpha modulation indices, when considering the grand average over all conditions (see Figure 2A) averaged over the previously defined time window of interest. These clusters constituted the regions of interests (ROIs) on which subsequent analysis was focused. Subsequently, in order to quantify individual hemispheric-specific bias with respect to modulation indices in the alpha range (MI(*α*)), we calculated the Hemispheric Lateralized Modulation (HLM) index per participant:

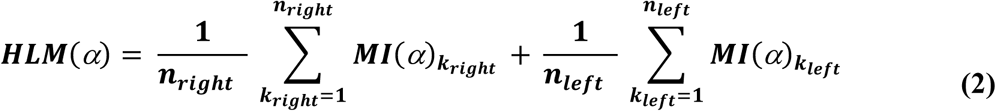

**Figure 2.**
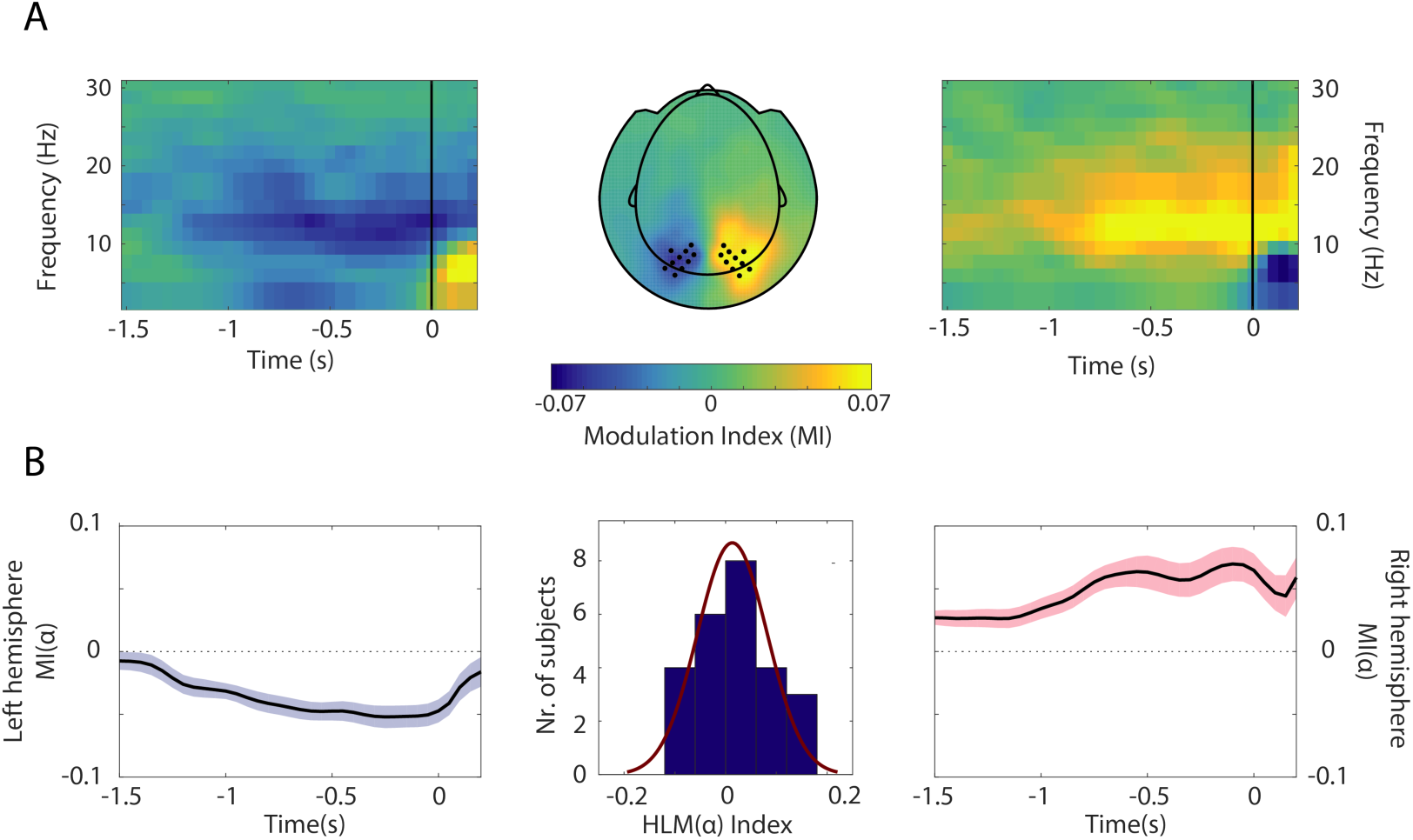
Grand average MI and HLM distribution across participants. (A) Time-frequency representations of power (TFRs) and topographical plot showing contrast between the ‘attention right’ – ‘attention left’ trials. A clear modulation is visible at posterior sensors in the alpha band (8 – 13Hz) in the –750 – 0ms interval (this time window being considered for the computation of HLM(*α*) indices in (B)). Sensors included in the left and right ROIs are marked as dots. Trials are locked to the onset of the contrast change (t = 0). (B) Side panels show the temporal evolution of modulation indices in the alpha range (MI(*α*)), averaged over sensors within left and right hemisphere ROIs. The magnitude (absolute value) of MI(*α*) progressively increased in the stimulus interval until the onset of the contrast change. Middle: distribution of HLM(*α*) indices across participants, computed over the ROIs and 8 – 13 Hz frequency band (see Materials and Methods). A normal density function is superimposed, denoting no hemispheric bias in lateralized modulation values across participants (Shapiro Wilk, *W* = .958, *p* = .392).

Where ***k_left_*** and ***k_right_*** denote sensors belonging to the aforementioned and previously defined left and right clusters, respectively. Please note that *MI*(*α*)*_k_* indices in Eq.2 (for both ***k****=*1, …, *n_left_* and ***k***=1, …, *n_right_*) are already a result of an average over timepoints of interest ***t***. Since MI(*α*) values were obtained by subtracting alpha power in ‘attend left’ trials from ‘attend right’ trials, and given that, as a result of attentional allocation, alpha power is suppressed in the hemisphere contralateral to the attended hemifield, a positive HLM(*α*) value indicated that a given participant displayed higher modulation of absolute magnitude of alpha power in the right compared to the left hemisphere, and vice versa (see Figure 2B). Analogously, lateralized indices (LI) of power modulation were computed for the alpha frequency band and for each subject at the cluster level according to:

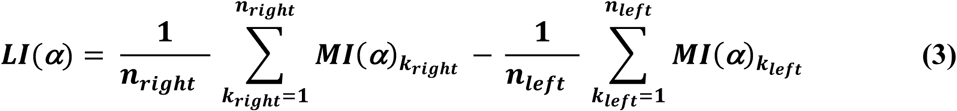

Also in this formula, ***k_left_*** and ***k_right_*** denote sensors belonging to the aforementioned left and right clusters, respectively. Since MI(*α*) values were obtained by contrasting alpha power in right versus left attention trials (see Eq.1), left hemisphere MI(*α*) were mostly represented by negative values, and right hemisphere MI(*α*) by positive values. Consequently, higher LI(*α*) indicated higher alpha lateralization for a given subject (i.e., higher interhemispheric difference in absolute alpha modulation).

### Structural data acquisition

T1-weighted images of three out of twenty-five participants were acquired on a 3 T MRI scanner (Magnetom TIM Trio, Siemens Healthcare, Erlangen, Germany), acquisition parameters: TR/TE= 2300/3.03 ms; FA=8°; FoV= 256 × 256 mm; slice thickness= 1 mm; Acquisition matrix= 0×256×256×0. For the remaining participants, a 1.5T MRI scanner was used (Magnetom AVANTO, Siemens Healthcare, Erlangen, Germany). Acquisition parameters: TR/TE= 2250/2.95 ms; FA=15°; FoV= 256 × 256 mm; slice thickness= 1 mm; Acquisition matrix= 0×256×256×0.

### Analysis

Structural analyses were conducted using the Integrated Registration and Segmentation Tool (FIRST) within FMRIB’s Software Library (FSL) v5.0.9 (www.fmrib.ox.ac.uk/fsl/, Oxford Centre for Functional MRI of the Brain, Oxford, UK). A standard 12 degrees of freedom affine registration to MNI152 space was applied to individual T1 images, adjusted with optimal sub-cortical weighting. Bayesian models implemented in the software are derived from a training based on previous manual segmentation of 336 datasets (provided by the Center for Morphometric Analysis (CMA, MGH, Boston) and applied to registered images to extract subcortical volumetric outputs for left and right hemispheres (see Figure 3A).

**Figure 3.**
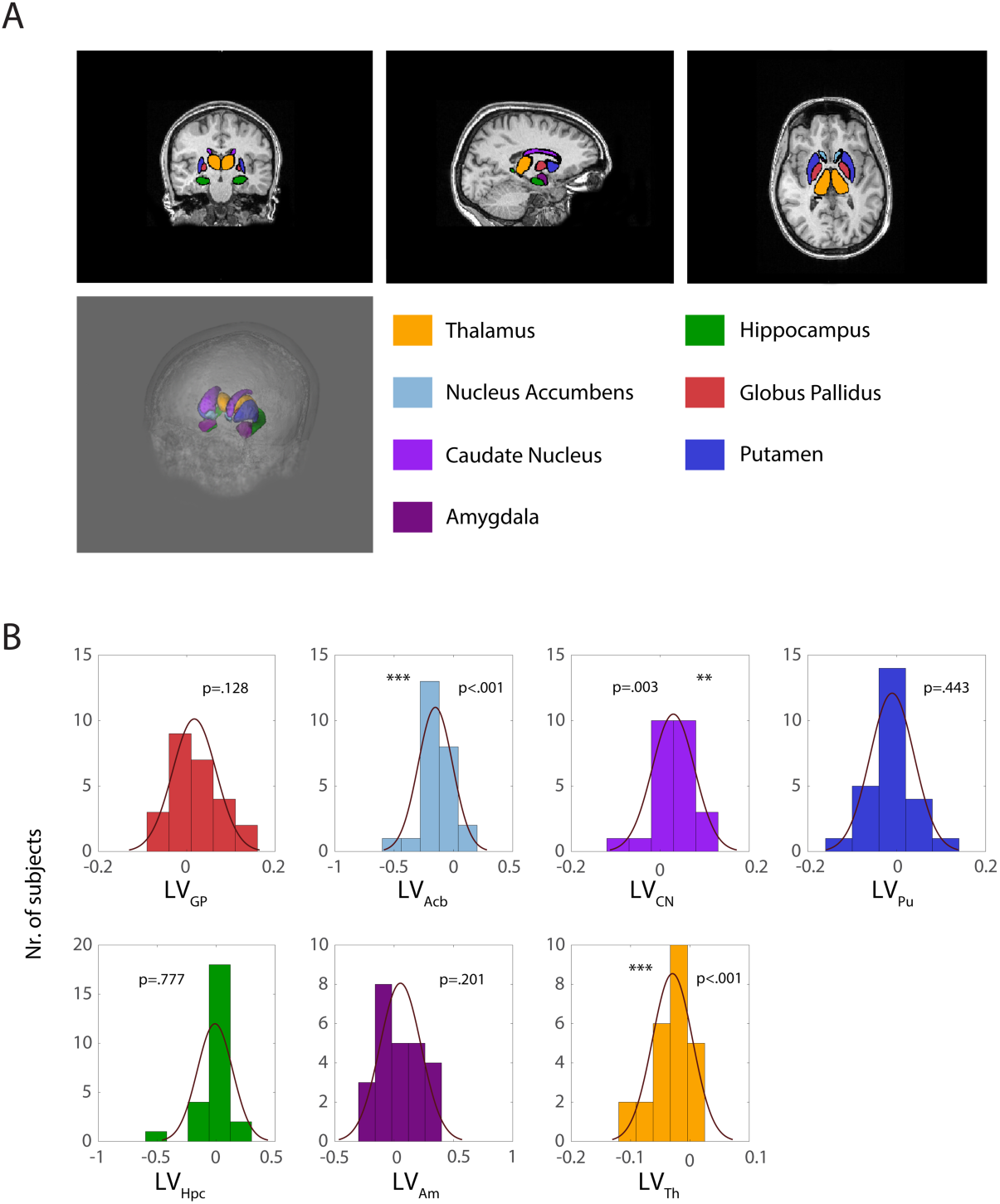
Basal Ganglia volumes resulting from semiautomated subcortical segmentation implemented. (A) Orthogonal view and 3D rendering. Subcortical volumes are overlaid as meshes on the anatomical MRI of one of the participants (following defacing procedure in Freesurfer, where voxels outside the brain mask with identifiable facial features were excluded (Bischoff-Grethe et al., 2007)). (B) Histograms with superimposition of normal density function, showing the distribution of subcortical lateralization indices for each substructure. In our sample, Acb and Th volumes were left lateralized (*p* = .0001 and *p*=.0003, respectively) while CN showed a right lateralization (*p* = .0029).

Given the reward components of the task we then focused on regions of the BG identified by the algorithm namely the Globus Pallidus (GP), Nucleus Accumbens (Acb), Caudate (CN), and Putamen (Pu), as well as the Thalamus (Th). Yet, an appropriate model attempting to describe basal ganglia influence on reward-related alpha modulation, needs to take into account the broader network of subcortical interconnections with neighboring nuclei. To this end, we included in the analysis the amygdala (Am) and the hippocampus (Hpc), whose interconnection has particularly been highlighted in the context of guided behavior when saliency processing was crucial (Paton et al., 2006; Zheng et al., 2017).

To compute hemispheric Lateralized Volume indices (*LV*) for each substructure of interest *s*, we used the following formula, which controls for individual differences in specific subcortical volumes via normalization by total bilateral volume, commonly employed to evaluate structural brain asymmetries (Guadalupe et al., 2016; Okada et al., 2016):

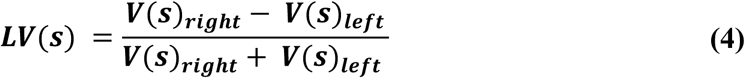

Where ***V_S_right__*** and ***V_S_left__*** represent respectively the anatomical right and left volumes (in voxels) for a given substructure *s*. Analogously to Eq.(2), a positive (or negative) LV_S_ index, in a given participant, indicated a greater right (or left) volume for a given substructure *s* (see Figure 3B).

### Statistical analysis

#### Generalized Linear Model

In order to determine the relationship between Basal Ganglia Lateralized Volumes (LV_S_) and electromagnetic indices (HLM(α)) we applied a generalized linear regression model (GLM), specifying subcortical volumes lateralization (LV_s_ values) as regressors and individual HLM values as the response vector.

To identify the optimal set of regressors to best predict HLM(α) indices, we pursued a model building strategy that would enable us to test the key hypothesis concerning the role of the BG and the thalamus. Hence, we considered all linear mixed-effects models including all possible combinations of at least 3 regressors (LV indices) using maximum likelihood estimation as parameter estimation method. We hence separately considered the models derived from all possible combinations of regressors, including either 2, 3, 4, 5 or 6 regressors (i.e. ROIs), by ‘picking’ the regressors from the lateralized subcortical volumes initially considered: (LV_GP_, LV_Acb_, LV_CN_, LV_Pu_, LV_Th_, LV_Am_, LV_Hpc_).

This resulted into a set of models for each of the four ‘options’ (2,3,4,5 or 6 number of regressors). Next, for each of the options, we derived the model associated with the lowest *Akaike Information Criterion* (AIC) and *Bayesian Information Criterion* (BIC), values commonly used for selection of best predictor subsets for a statistical model. Upon selection, we ended up with the four ‘best’ models, representative of each of the four options described above.

The final step, was to identify the ‘winning model’ among the selected ones (i.e. lowest AIC and BIC values) and compare it with the full model (7 regressors), which included the whole set of substructures, according to the formula:

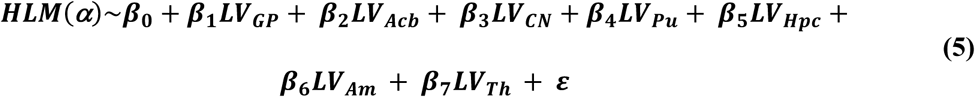

All subsequent analysis on the relationship between volumetric and oscillatory data specifically focused only on the subcortical structure(s) associated with a significant β coefficient in the model in Eq.(5), below referred as LV_s_.

#### Cluster based permutation test

To evaluate whether the linear association between LV_s_ and HLM was effectively limited to the alpha band, a cluster based permutation approach (Maris and Oostenveld, 2007) was employed over the full time frequency spectrum of interest. This method effectively allows to statistically control for multiple comparisons over all time and frequency points of interest. After selecting the *a-priori* sensors belonging to the formerly specified ROIs, we considered a permutation distribution of regression coefficients derived from randomly pairing participants’ LV_s_ value (independent variable) and modulation indices (MI(*f*)) 1000 times. At every time-by-frequency point, the actual regression coefficient was evaluated against the aforementioned distribution by means of a specified critical α value. Afterwards, a time-frequency map of the cluster level statistics was derived showing sets of sensors associated with a significant effect.

An equivalent approach was later applied to investigate possible hemisphere-specific differences in alpha modulation between participants showing a right or left lateralized substructure s. Directionality of lateralization was determined by median split of the distribution of LV_s_ per participant, producing two subgroups of N=12, representing subjects with a larger left or right volume of substructure s. After having a-priori averaged across the time-frequency spectrum of interest ([-1500 - 0] ms, 8-13Hz), MI(α) values at every sensor were compared between the two subgroups (right vs left lateralized substructure). The actual t-value was then compared with a permutation distribution of t-statistic derived from randomly partitioning indices between the two groups 1000 times. As a result, a topography map was plotted displaying eventual cluster(s) of sensors associated with a significant t-value (i.e. a significant difference in MI(α) between subgroups).

#### Comparison between Pearson’s correlation coefficients

Finally, we aimed at comparing the association between the derived structural and functional lateralization indices in different value salience occurrences. To this end, we calculated HLM(*α*) values for each participant separately for the three reward-related contingencies and computed the Pearson’s correlations with LV_S_ indices which displayed a significant β as arising from the model in Eq.(5). We statistically assessed the difference in correlation coefficients between the three experimental conditions considered, according to the method described in (Wilcox, 2016a). The test implements a percentile resampling technique by generating a bootstrap sample of the difference of the correlation coefficients between the overlapping variable LV_GP_ (Y) and the two variables representing the HLM(*α*) for the two experimental conditions (VO levels) to be compared (X_1_, X_2_). As suggested in the method, we used a Winsorized correlation to achieve a robust measure of association between variables. This transformation has been shown to effectively control for the influence of outliers on the correlation estimate (Wilcox, 2016b). A confidence interval was then computed on the resulting bootstrap distribution, to assess the statistical significance of the actual difference between correlation coefficients describing the different VOs.

#### Behavioural data analysis

To assess whether subjects displayed a spatial bias during the task, we first averaged across left and right cued trials separately, averaged across all conditions (i.e., irrespective of value-saliency occurrences (VO)). We then employed paired t-test on the derived reaction times (RT) and accuracy (ACC) (expressed as percentage of correct responses) measures for the left and right cued trials. Secondly, we divided trials according to VO pairings, averaging left and right cued trials, to determine whether behavioural performance varied as a function of saliency in both RT and ACC. We here employed one-way repeated measures ANOVA to assess whether group means in the three conditions significantly differ from each other. We also considered individual lateralized measures of RT and ACC across different VO conditions. To this end, behavioural asymmetries in performance (BA) for both measures were calculated according to:

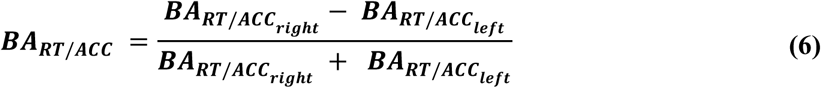

Where BA_RT *right*_ and BA_RT *left*_ represent mean reaction times for ‘attend right’ and ‘attend left’ trials, respectively. A positive BA_RT_ for a given subject indicated faster responses when a participant was validly cued to the left compared to the right hemisphere, while negative values indicated the opposite pattern. Consequently, positive BA_ACC_ values indicated higher accuracy on ‘attend right’ trials compared to ‘left attend’ trials, and vice versa.

A one-way repeated measures ANOVA was employed to test the difference across group means in the three VO conditions examined.

In a next step, we sought to investigate the possible association of behavioural performance with structural and functional hemispheric lateralization, we used Pearson’s correlation to examine the association of individual asymmetries in accuracy (BA_ACC_) and reaction times (BA_RT_) with individual HLM(*α*) and LV values of subcortical structures which showed significant correlation with HLM(*α*).

In a last step, we employed a general linear model (GLM) in order to assess whether spatial biases in behavioural performance could be explained by a combination of the other variables, namely HLM(*α*) and the LV indices of the subcortical areas considered, according to the formula:

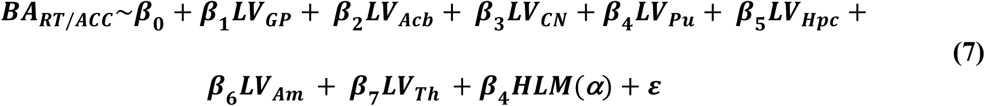

## Results

We acquired structural and electrophysiological data from 25 participants. Participants’ performance was tested during a covert attention paradigm, where Chinese symbols served as targets and distractors (Figure 1). During a learning phase, prior to the actual task, the stimuli were associated with different values (positive, negative or neutral). In the testing phase, a central cue probed an upcoming contrast variation of the target, which appeared either at 1450 or 2350 ms, predicting its position in 95% of the trials. Participants were instructed to indicate, with button press, the direction of the contrast change, which could either increase or decrease with equal probability. MEG data, eye-tracking and behavioural responses were acquired during the testing phase. Time-frequency representations of power were calculated from MEG trials after preprocessing and artifacts rejection. Power modulation (MI) indices were computed by contrasting power in trials where participants were validly cued to the right (*attend right* trials) with trials where participants were validly cued to the left (*attend left* trials) (see Eq.(1), *Materials and Methods*).

As presented in the previously reported results (Marshall et al., 2017), we confirmed that participants displayed a clear modulation of alpha band activity in parieto-occipital sensors (MI(*α*)): when covertly orienting attention to the cued side, alpha power decreased in the contralateral hemisphere while it increased relatively in the ipsilateral hemisphere (Figure 2A). The magnitude of alpha power modulation, as reflected by MI(*α*), progressively increased until the target changed contrast (Figure 2B). To best quantify the modulation, we focused our analysis on the 750 ms interval immediately preceding the onset of the contrast change. Next, right and left ROIs were identified as clusters of symmetric pairs of sensors showing the highest alpha lateralization values (see *Materials and Methods*) (i.e., sensors displaying highest interhemispheric difference in alpha modulation).

Starting from the assumption that, to a certain extent, an inter-subject variability in the ability to modulate alpha power – in absolute value – must exist in the right compared to the left hemisphere (and vice versa), we sought to quantify individual hemispheric biases in the ability to modulate alpha activity. To this purpose, hemispheric lateralized modulation of alpha power (HLM(*α*)) values were then computed for each participant by summing the average MI(*α*) in the right and left hemisphere ROIs (see Eq.(2), *Materials and Methods*). As a result of this computation, positive HLM(*α*) values would demonstrate that a given subject was better at modulating their right, compared to left, hemisphere alpha power, while a negative index would reflect higher ability to modulate alpha power on their left, compared to right, hemisphere. The histogram in Figure 2B depicts the distribution of hemispheric biases related to attentional modulation of alpha power. HLM(*α*) indices ranged from about −0.1 to 0.1 (i.e. a 20% variation) but they were normally distributed around zero across participants (Shapiro Wilk, *W* = .958, *p* = .392).

### Volumetric asymmetry of basal ganglia in relation to hemispheric lateralized alpha modulation

The next step was to determine whether the biases in the ability to modulate left versus right hemisphere alpha (HLM(*α*)) was related to individual hemispheric lateralization of subcortical structures. A semi-automated segmentation tool implemented in FMRIB’s Software Library (FSL), was used to estimate volumes for the left and right subcortical and limbic structures, namely: Globus Pallidus (GP), Nucleus Accumbens (Acb), Caudate Nucleus (CN), Putamen (Pu), Hippocampus (Hpc), Amygdala (Am) and Thalamus (Th). We then calculated the hemispheric lateralized volumes (LV) for each set of structures (see *Materials and Methods*, Eq.(4)). Positive (or negative) LV_s_ values, for a given participant, indicated whether a specific substructure *s* was larger in the right compared to the left hemisphere (and vice versa). Further analysis revealed that, over subjects, the Acb and Th were significantly left lateralized (*z* = −3.78, *p* = 1.56*×*10^−4^ and *z* = −3.59, *p* = 3.28*×*10^−4^, respectively; two-sided Wilcoxon signed rank test), whilst the CN was right lateralized (*z* = 2.97, p = .003). For the other substructures, no significant lateralizations were identified.

In order to corroborate that the observed anatomical lateralizations were representative of the population and not merely a fluke in the dataset, we applied the same analysis to a pool of 612 independent, anonymized anatomical MRI scans internally available at the Donders Institute. We hence estimated left and right volumes for the same subcortical structures considered in our study, and derived respective LV indices (see Eq.(4)). Importantly, the same direction of lateralization in all the substructures was found in the Donders dataset and the one reported in our sample. Specifically, Th and the Acb were significantly left lateralized (*z* = −14.0, *p* = 1.4*×*10^−44^ and *z*= −17.04, *p* = 4.1*×*10^−65^, respectively), whilst the CN was right lateralized (*z* = 13.0, *p* = 1.2*×*10^−38^). In addition, the GP was found to be right lateralized (*z* = 4.0, *p* = 5.9*×*10^−5^, an effect only observed as trend in our dataset), as well as the Hpc (*z* = 4.95, *p* = 7.3*×*10^−7^), while the Pu was left lateralized (*z* = −3.27, p = .001). These surprising significant lateralization biases in a large dataset are highly interesting, given they suggest differential lateralizations of subcortical structures on a population level. Moreover, they support the conclusions drawn in the context of the study.

Given the volumetric variability in the set of substructures considered for the segmentation protocol, we performed a cross-correlation analysis between the different substructures, including left and right volumes, to query about a potential bias in the segmentation algorithm. No significant effects were found (positive or negative correlations, all *p-values* >0.688; highest negative correlation *R*=-.084) indicating that, if a given structure is larger for a given subject, this doesn’t imply a bias in the segmentation protocol (e.g. to the expense of neighbouring, allegedly smaller areas).

To investigate whether individual subcortical asymmetries (LVs, as defined in Eq.(4)) predicted differences in hemispheric lateralized modulation of alpha power (HLM(*α*)), we implemented a GLM, where LV indices were included as multiple explanatory variables for the response variable (individual HLM(*α*)).

As described in *Materials and Methods* section, we pursued a model building strategy that would enable us to test the key hypothesis concerning the role of the BG and the thalamus. We analyzed all linear mixed-effects models derived from all possible combinations of at least 2 regressors (LV indices). Following this evaluation, we identified the 5 regressors model as the best (with AIC= −71.23 and BIC=−62.70). This model included as regressors LV_GP_ (*p*=4*×*10^−4^), LV_Acb_,(*p*=.036) LV_Pu_ (*p*=.028), LV_Hpc_ (_p_=.144) and LV_Th_ (*p*=.022).

The selected 5 regressors model provided a better estimation of HLM(*α*) given the set of predictors, when compared to the full model (including all the 7 substructures), which had AIC of −67.5 and BIC=-56.5 (see Eq.(5)).

Despite having defined the 5 regressors model as the optimal set for predicting HLM(*α*), we proceeded our analysis with the full model, describing the predictive value of the whole set of lateralized subcortical volumes. The underlying aim was to be more conservative, address potentially confounding effects of neighboring regions, and to include the full set of BG structures for a complete overview of their effects. We therefore report related results below. Of importance here is to note that, in the 5 regressors *winning* model, based on the output of our model comparison, the results concerning our structures of interest still held: LV_GP_ and LV_Th_ significantly predicted HLM(*α*) values.

The full model was associated with a significant regression: when considering the grand average of all conditions, a linear combination of all the subcortical LVs was able to explain HLM(*α*) values (*F*_7,17_ = 3.37, *p* = .019, adjusted *R*^2^ = .409). When assessing each predictor individually, only the beta coefficients for LV_GP_ and LV_Th_ were found to be significantly higher than zero (partial correlation: *p* = .004 and *p* = .028) (Figure 4A). Hence, when controlling for the other explanatory variables in the model, only GP and TH asymmetry (LV_GP,_ LV_TH_) significantly contributed to explain biases in hemispheric lateralized alpha band modulation (*β* = 1.768 and *β* = 1.924, respectively). The independent contribution of GP and TH lateralization is visible in Figure 4B,C showing the partial regression plots for LV_GP_ and LV_TH_ in relation to the HLM(*α*) values. We conclude that hemispheric biases in GP and TH volume are predictive of the individual abilities to modulate left versus right hemisphere alpha. Precisely, subjects presenting a larger GP volume in the left hemisphere compared to the right, also displayed a higher ability to modulate alpha power (in absolute value) in the left visual hemisphere compared to the right (and vice versa); the same association holding for the Th in relation to HLM(*α*) values.

**Figure 4.**
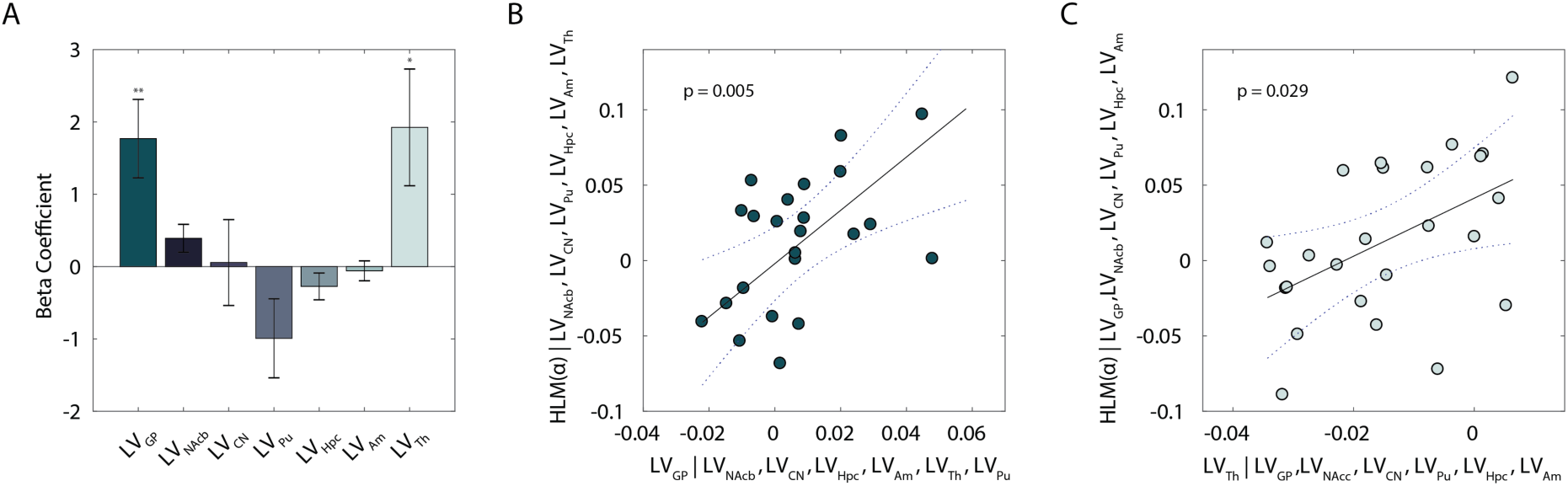
Lateralization of individual subcortical structures in relation to alpha hemispheric lateralized modulation (HLM) in the task. (A) Bar plot displays the Beta coefficients associated with a general linear model where LV values were defined as explanatory variables for HLM(*α*). Error bars indicate standard error of the mean. Asterisks denote statistical significance; \*\**p* < .01. (B) Partial regression plot showing the association between LVGP and HLM(*α*), while controlling for the other regressors in the model in (A). (C) Partial regression plot showing the association between LVTh and HLM(*α*), while controlling for the other regressors in the model in (A). Given Eqs.(1) and (2) (see *Materials and Methods*), positive HLM(*α*) values indicate stronger modulation of alpha power in the right compared to the left hemisphere, and vice versa; similarly, positive (or negative) LVS indices denote greater right(or left) volume for a given substructure *s*. The dotted curves in (B) and (C) indicate 95% confidence bounds for the regression line, fitted on the plot in black.

### Hemispheric asymmetry of globus pallidus correlates selectively with power modulation in the alpha band

In order to better interpret the GLM results, we assessed whether the linear relationship arising from the model was restricted to the alpha band: to this end, a non-parametric approach was implemented to further explore the LV_GP_ and LV_TH_ in relation to HLM(*α*). This method allows circumvention of the multiple comparison problem over frequency and time points by evaluating the full low-frequency spectrum (2-30 Hz) from −1500 to 0ms (Maris and Oostenveld, 2007). We therefore conducted a cluster-based permutation test using a dependent samples regression *t*-statistic to evaluate the effect (linear association between LV_GP/TH_ indices and HLM over all frequencies) at the sample level. A *p*-value of .05 was chosen for thresholding the *t*-statistic of the permutation distribution and a critical value corresponding to alpha = .025 (two tailed) was considered for the cluster-level regression test statistic. As depicted in Figure 5A, we observed a significant cluster (*p* = .008) extending for 1000 ms window prior to contrast change (i.e., when covert attention was deployed to the cued stimulus), which confirmed a positive linear association (positive *t*-value) between LV_GP_ asymmetry and hemispheric lateralized modulation (HLM) of power constrained to the alpha frequency range. When applying the same analysis to the Th asymmetry in relation to HLM, no significant clusters of sensors were identified.

**Figure 5.**
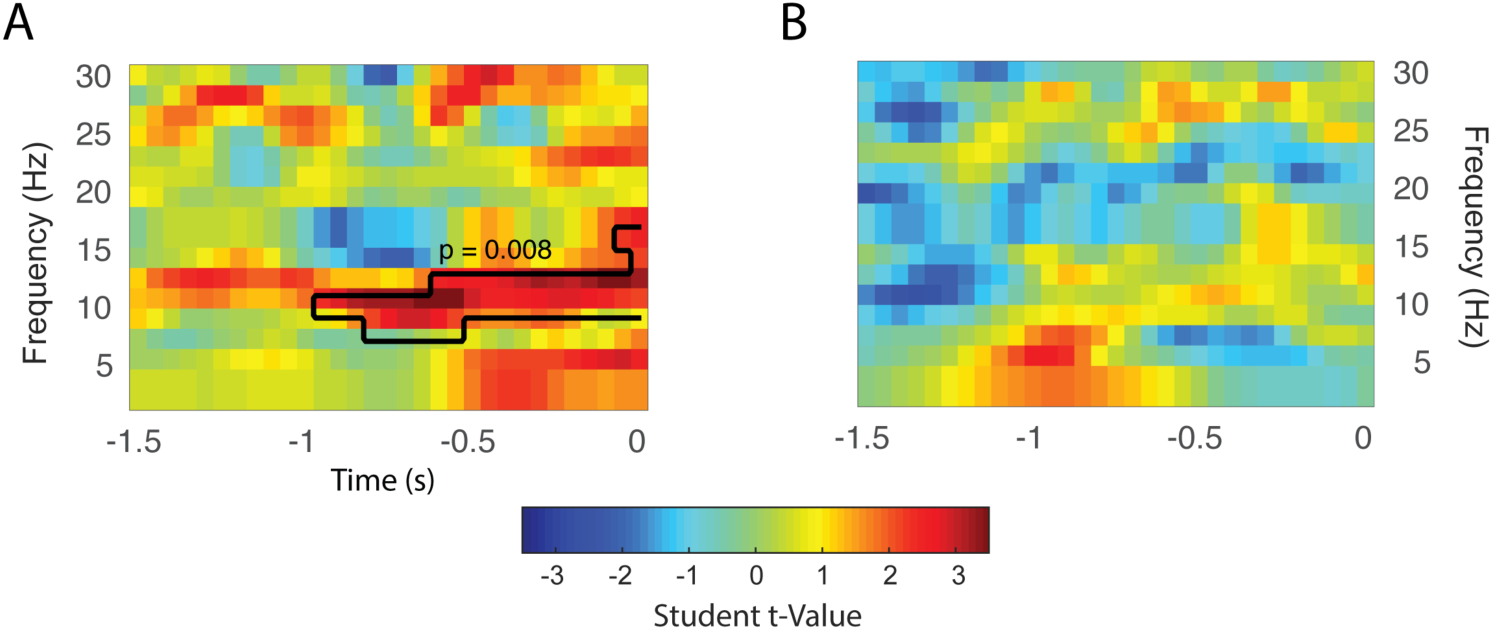
Time-frequency representation of regression coefficient t-statistics on the linear relationship between low frequency power modulation (MI(*f*)) and LVGP (A) and LVTh (B) indices, averaged over ROIs (see Fig. 2A). A black outline is used to highlight the significant time-frequency cluster found. For the LVGP, the analysis revealed a clear α-band-limited association between the variables across the full time-window of interest (see *Materials and Methods*) extending up to 1s prior to the response.

### Hemisphere specific relations between alpha modulation and GP asymmetry

Given the specific association found between LV_GP_ and HLM(*α*), arising from the previous analysis, we decided to further investigate the hemisphere-specific influence of GP volumetric asymmetry on alpha modulation indices. For this purpose, we sought to compare average left and right hemisphere MI(*α*)s of participants according to the direction of GP lateralization. This was done by means of median split of the LV_GP_ distribution, hence resulting in two subgroups, that either had a bias towards a larger left than right GP volume or vice versa (see *Materials and Methods*). Figure 6 displays a topographical representation of MI(*α*) values per subgroup (A), together with individual raw data points, superimposed on bars representing average values per ROIs per each subgroup (B) and distribution of individual HLM(*α*) values (C). Consistent with the GLM results, participants with a larger right than left GP, also displayed a higher modulation of alpha band (in absolute value) in the right hemisphere compared to the left. Given that the assumption of normality, required to perform a mixed-effect ANOVA, was not met for the distribution of MI(*α*) indices in the two subgroups, we implemented a non-parametric cluster-based permutation test to compare the MI(*α*) between the two aforementioned subgroups (averaged across specific time and frequency band of interest), employing an independent sample t-test score, and then comparing it with the resulting permutation distribution. This allowed us to explore whether there was a hemispheric-specific difference in the two subgroups in the extent of absolute alpha modulation. The test indicated a significant cluster of sensors over right posterior channels (p= .027), hence including the previously defined right ROI and denoting a significant difference in the right hemisphere absolute alpha modulation (MI(*α*)) between the two subgroups. These results might suggest that the linear association arising from the GLM (Figure 4A, B) in relation to the association between LV_GP_ and HLM(*α*), was largely driven by right hemisphere alpha modulation. The analogous analysis was conducted on the median split of the distribution of LV_TH_ indices. In this case we did not find interhemispheric dominance in alpha modulation indices related to lateralization of the thalamus in the right compared to left hemisphere.

**Figure 6.**
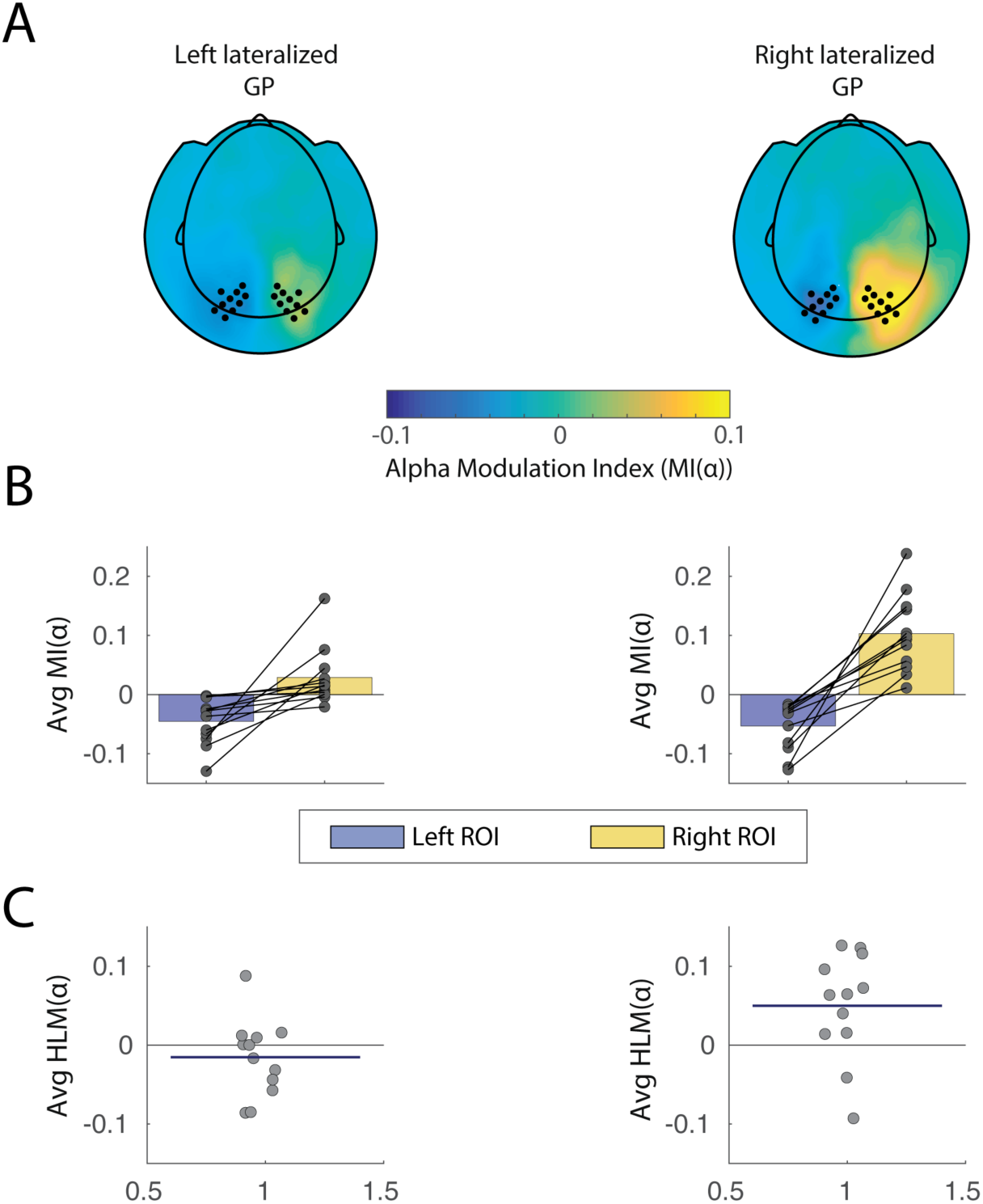
Alpha modulation indices for left and right hemispheres associated with two subgroups of the sample. (A) Topographical plot of MI(α) values for the two participants groups, clustered according to directionality of GP lateralization (right vs left lateralized GP). Left and right sensors of interest are marked as dots and correspond to the same ROIs as in Figure 2. (B) Individual datapoints superimposed on bar graph showing individual scores and MI(α) averaged over ROIs in the two subgroups. As indicated in the cluster-based permutation results, a difference is particularly observable for right hemisphere alpha modulation between the two groups, being higher in participants exhibiting a right lateralized GP. (C) Individual datapoints showing HLM(α) scores for all participants. The horizontal blue line superimposed on the data indicates average HLM(α) index for each subgroup

### The involvement of globus pallidus and thalamus in relation to stimulus-value associations

Crucially, we aimed at assessing whether the level of value-saliency occurrences (VO) in a given trial influenced the association between the structural and functional lateralization indices arisen from the GLM. We first calculated HLM(*α*) (see Eq.(2), *Materials and Methods*) values for each participant, separately for the three VO levels, namely two, one and zero value saliency occurrences (see *Materials and Methods*). We then examined Pearson correlations between HLM(*α*) and LV values for both GP and Th, which showed a positive significant β in the model, across the three levels considered (Figure 7, Figure 8). LV_GP_ significantly correlated with HLM(*α*) only in trials where both target and distractors had value-salience (two VO) (*r* = .68, *p*=1.75*×*10^−4^; Figure 7A). This denotes that, in trials with two value-salient items presented, participants exhibiting a right lateralized GP volume, also displayed a stronger alpha modulation in the right compared to the left hemisphere, and vice versa. LV_GP_ did not significantly correlate with HLM(*α*) when only one or none of the stimuli presented were associated with a salient value (*p* = .144 and *p* = .314, respectively; Figure 7A).

**Figure 7.**
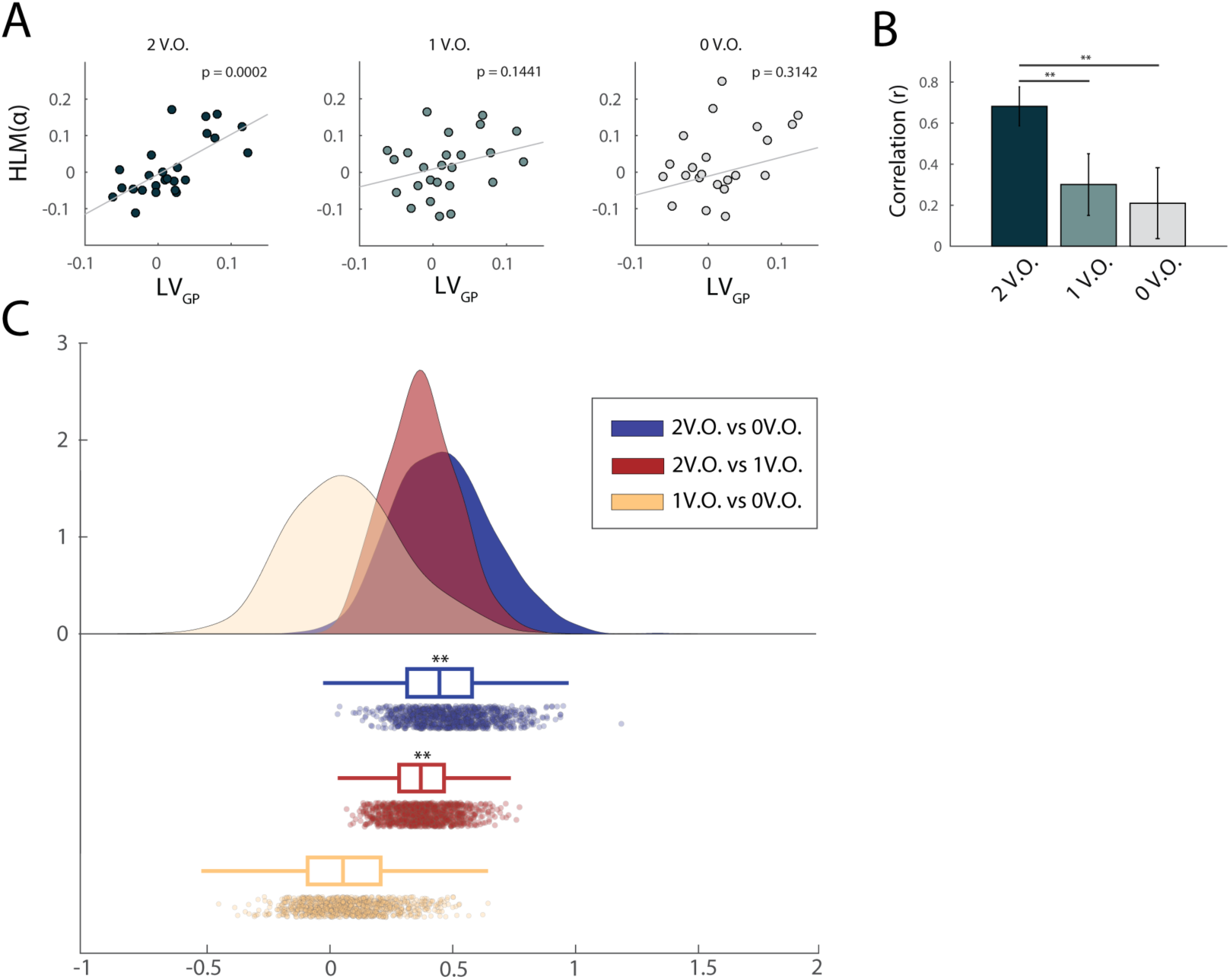
Linear association between GP volumetric asymmetry and alpha modulation asymmetry as a function of value-saliency occurrences in the task. (A) Correlation between GP volume lateralization and HLM(α), grouped accordingly to the number of value-salient stimuli in the trials (see *Materials and Methods*). From left to right, respectively, two, one and zero value-saliency occurrences are displayed. GP asymmetry significantly explained HLM(α) only when value-salient stimuli featured as both target and distractors, irrespective of their valence (*r* = .68, significant at the *p* < .001 level after Bonferroni correction for three comparisons). (B) The association between HLM(α) and GP volume lateralization increased as a function of value saliency in the task: the linear relationship was stronger when two value-salient stimuli were presented, when compared to conditions characterized by either one or value-salience pairings (95% CI [.106, .672] and [.125, .897], respectively for the two comparisons). This suggests that, when both target and distractor were associated with a salient value, participants exhibiting bigger GP volume in the left hemisphere than in the right hemisphere, were also better at modulating alpha oscillations in the left compared to the right hemisphere. Asterisks denote statistical significance; \*\**p* < .01. (C) Raincloud plot (Allen et al., 2018) showing the bootstrap distribution of the difference in pairwise correlation coefficients examined.

**Figure 8.**
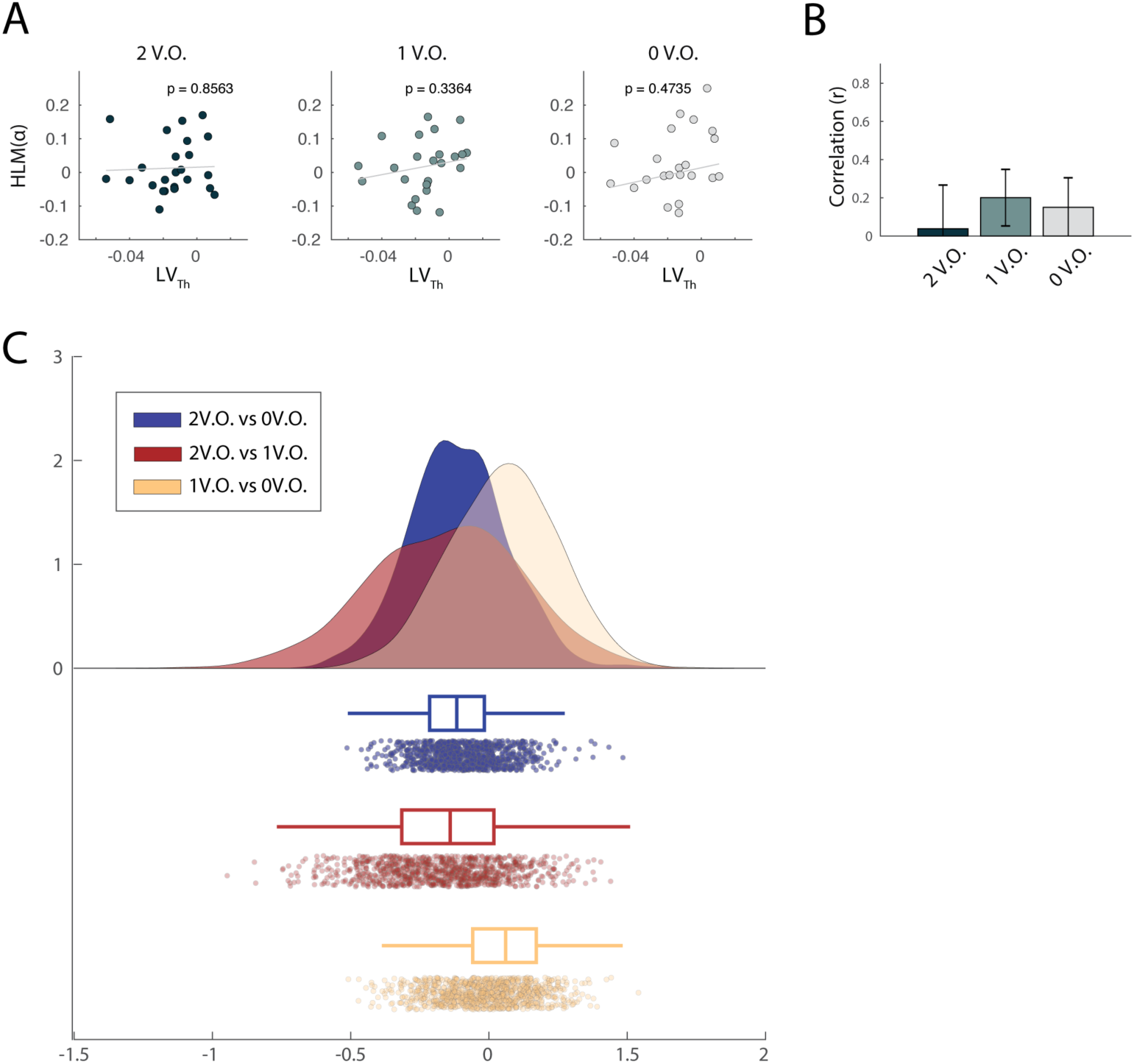
Linear association between Th volumetric asymmetry and alpha modulation asymmetry as a function of value-saliency occurrences in the task. (A) Correlation between TH volume lateralization and HLM(α), grouped accordingly to the number of value-salient stimuli in the trials (see *Materials and Methods*). From left to right, respectively, two, one and zero value-saliency occurrences are displayed. When considering individual correlations between Th asymmetry and HLM(α), no significant linear relationship was found. (B) The association between the two measures also didn’t significantly differ as a function of saliency in the trials. (C) Raincloud plot showing the bootstrap distribution of the difference in pairwise correlation coefficients examined.

In order to statistically quantify the influence of the stimulus-value association on the relationship between LV_GP_ and HLM(*α*), we compared *robust correlations* in the three conditions according to the bootstrap method described in (Wilcox, 2016c) for dependent overlapping correlations (see *Materials and Methods*). The correlation between LV_GP_ and HLM(*α*) in trials with two occurrences of value-salience, significantly differed both from the condition characterized by one (95% CI [.106, .672]) and zero occurrences (95% CI [.125, .897]). This confirmed that the association between lateralized GP volume and alpha modulation bias significantly increased as a function of the number of value-salient occurrences in the task (Figure 7B). Bootstrap distributions of the pairwise difference in correlation coefficients is shown in Figure 7C. We performed the same analysis in order to assess whether value-saliency occurrences mediated also the association between LV_TH_ and HLM(*α*). When considering the correlation indices in the three conditions separately, no significant linear relationship was found between the two indices (Figure 8A). Also in this case, when comparing robust correlations between the three conditions, according to the same method above, no significant difference was found (Figure 8B,C). This suggested that the relationship between thalamus volumetric lateralization and alpha modulation arising from the model in Eq.(5), was not driven by the number of value-salient occurrences in the task.

### Behavioural analysis

#### Stronger alpha lateralization is associated with better behavioural performance in the task

At the behavioural level, we expected to corroborate existing literature linking alpha oscillations to behavioural performance in spatial attention tasks. In order to disentangle possible confounds derived from the value component of the task, we first considered only neutral trials (holding 0 V.O.). We then performed a trial-based analysis, by first grouping, for each subject, fast and slow trials, with respect to the reaction times distribution median. This was done separately for left-cued and right-cued (valid) trials. For each subject, we then computed alpha modulation indices (MI(α)) of derived fast and slow trials, according to Eq.1 (see Materials and Methods). Next, we averaged MI(α)s in slow and fast trials across subjects. Figure 9A shows the topographical representation of MI(α)s values for the two trial groups (fast versus slow trials). Figure 9B shows individual and mean values for MI(α), over left and right ROIs in the two subgroups, while in panel C are displayed individual and averaged LI(α)s per subgroup. To statistically assess the difference in alpha lateralization between the two subgroups, we compared LI(α)s (according to Eq.(3)) between fast and slow trials, by means of dependent sample t-test. This revealed that, on average, subjects displayed a stronger alpha lateralization in fast trials, as compared to slow trials (t_(24)=_2.27, p=.032), when no saliency processing was required (0 V.O. trials).

**Figure 9.**
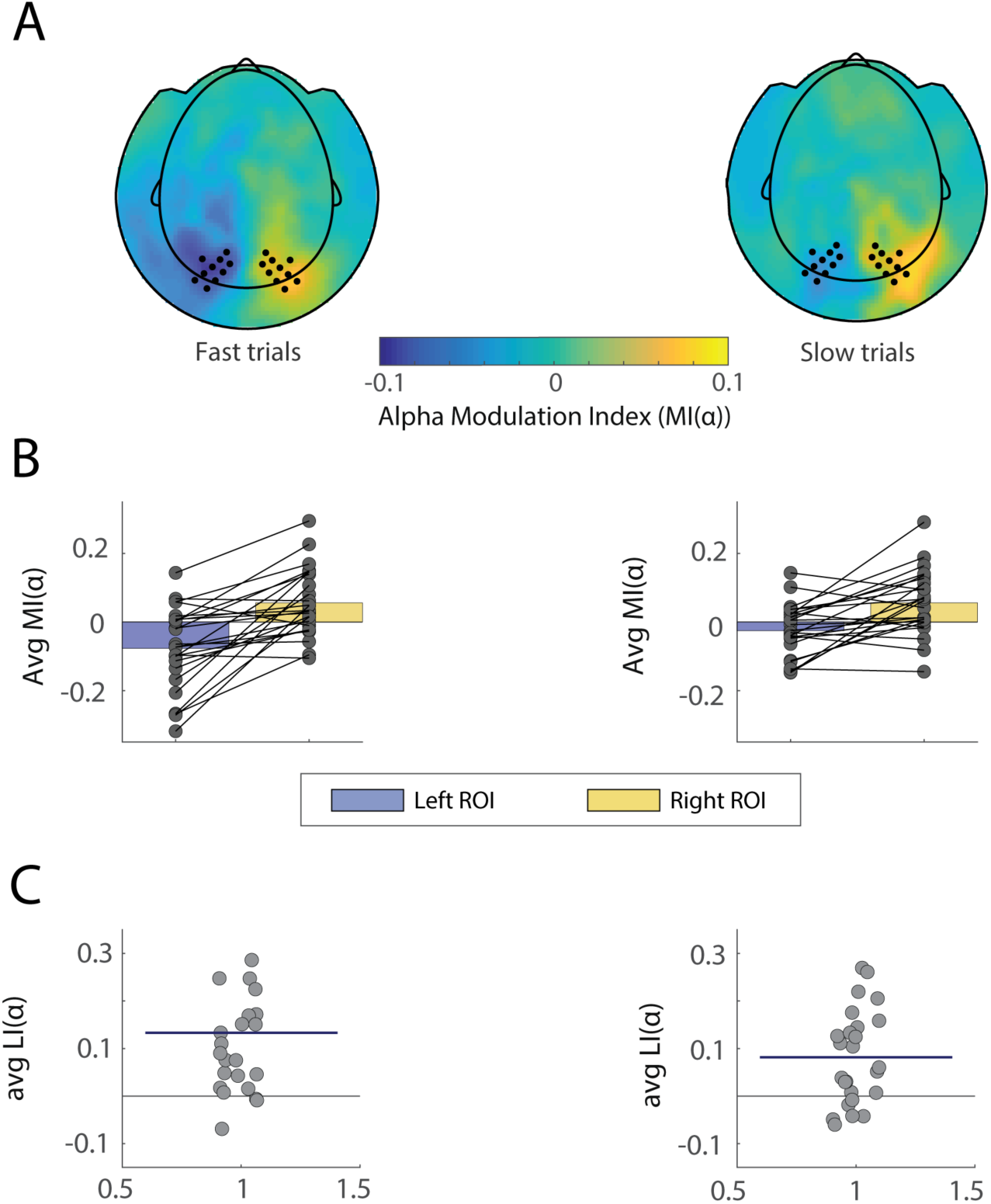
Alpha modulation indices for left and right hemispheres associated with fast versus slow trials, neutral condition only. (A) Topographical plot of MI(α) values for the two trial groups, clustered according to median split of reaction times (fast versus slow trials). Left and right sensors of interest are marked as dots and correspond to the same ROIs as in Figure 2. (B) Individual datapoints superimposed on bar graph showing individual scores and MI(α) averaged over ROIs in the two subgroups. (C) Individual datapoints showing LI(α) scores for all participants (difference in MI(α) values between right and left ROIs above). The horizontal blue line superimposed on the data indicates average LI(α) index for each subgroup.

To be able to generalize the effect to the whole task, we performed the same trial-based analysis described above on all the conditions irrespective of their V.O. levels. The analysis showed that, overall, subjects produced a significantly stronger alpha lateralization in fast trials, as compared to slow trials, irrespective of trial type (t_(24)_=2.63, p=.014).

#### Behavioural performance is not dependent on saliency occurrences in the task

At the behavioural level, we sought to investigate whether subjects displayed a spatial bias in task performance, irrespective of the value-saliency levels. To this end we performed a paired t-test to assess whether participants’ performance differed between left and right cued trials, in both reaction times (RT) and accuracy measures. No behavioural spatial bias was found neither in RT (p=.341) nor in accuracy (p=.572) values. Secondly, we examined whether value-salient occurrences (VO) levels, modulated participants’ behavioural performance. We then compared mean RT and accuracy for the three VOs levels (see *Materials and Methods*). There were no statistically significant differences between the three groups, as determined by one-way ANOVA, in mean RT (F_(2,72)_ =.004, *p*=.995) (Figure 10A) nor in mean accuracy (F_(2,72)_=.003, *p*=.996) (Figure 10C). We then tested whether a behavioural spatial bias occurred across value saliency occurrences (i.e., whether subjects displayed a difference in RT or accuracy asymmetry across VOs). We computed measures of behavioural asymmetry in accuracy (BA_ACC_) and reaction times (BA_RT_) (see Eq.(6), *Materials and Methods*). Analogously to the method used to compute HLM(*α*), we created asymmetry indices for every subject by contrasting behavioural measures for *attend right* with *attend left* trials. As such, a positive BA_RT_ would indicate that subjects were faster when cued to the left compared to the right hemisphere, and vice versa. Similarly, positive BA_ACC_ indices reflected higher accuracy when cued to the right compared to the left hemisphere. With the method aforementioned, we performed a one-way ANOVA to assess whether a significant difference in behavioural bias occurred across the three VO conditions. Neither BA_RT_ nor BA_ACC_ values significantly differed across VOs (F_(2,72)_=.191, *p*=.826 and F_(2,72)_=.669, *p*=.515, respectively) (Figure 10B, 10D).

**Figure 10.**
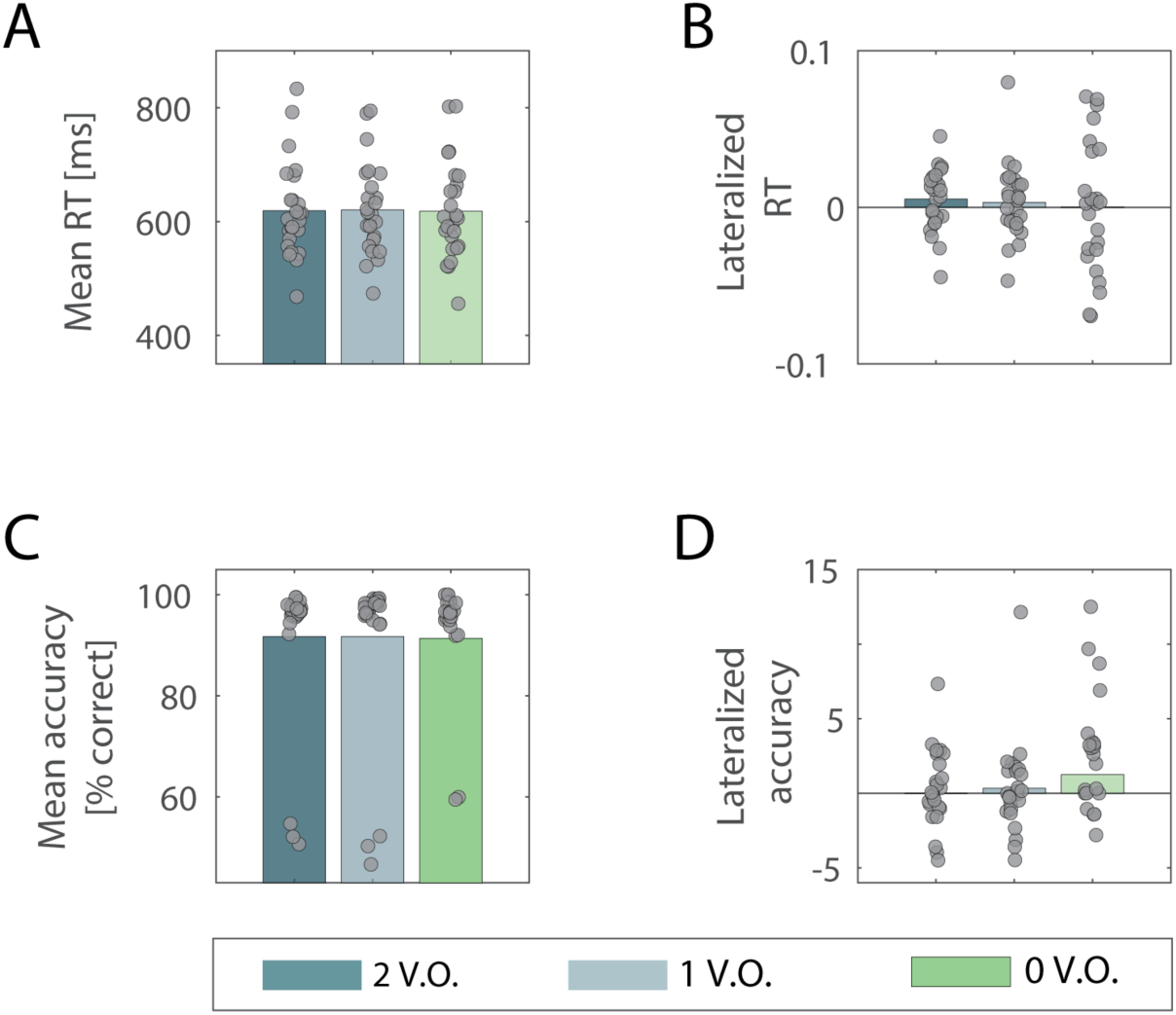
Mean and lateralized reaction times (RT) and accuracy values across the three value-saliency occurrences in the task. Mean RT (A) and accuracy (C) values averaged across participants in the three value-salient occurrences conditions in the task. No significant difference was found between groups by means of one-way repeated measures ANOVA, indicating that different levels of value-saliency pairings didn’t influence behavioural performance. No significant difference emerged also when comparing average lateralized values of RT (B) and accuracy (D) across the same conditions, and by means of same statistical analysis, indicating that the behavioural spatial bias was not affected by the different levels of value-saliency pairings. Respective individual scores are superimposed on bars in all plots.

The resultant lack of a relationship between spatial bias in task performance and degree of saliency processing required (V.O.s) is likely explained by the orthogonalization of attentional orienting and stimulus-value associations in the task.

With the aim of determining a potential link between lateralized indices of behavioural performance and the anatomical (LVs) and functional (HLM(*α*)) lateralization indices of interest, we employed three separate GLMs to assess whether a linear combination of BA_RT_ and BA_ACC_ values could explain LV_GP_, LV_TH_ and/or HLM(*α*) indices. Neither LV_GP_ nor LV_TH_ could be explained by the behavioural lateralized measures (*F*_1,23_=.18, *p*=.834, adjusted *R^2^*=-.07 and *F*_1,23_=.16, *p*=.849, adjusted *R^2^*=-.07). The same result held for the prediction of HLM(*α*), yielding also in this case no significant regression coefficients (*F*_1,23_=1.17, *p*=.33, adjusted *R^2^*=-.01).

Last, we investigated whether individual behavioural spatial biases could be accounted for by a combination of the other measures examined. To this end, we considered all subcortical LV_S_ and HLM(*α*) indices and specified them as regressors in a general linear model (see Eq.(7), *Materials and Methods*), in order to determine whether they could explain biases in RT and accuracy (BA_RT_ and BA_ACC_). No significant regression was found which could account for BA_RT_ indices (*F*_8,16_=.85, *p*=.570, adjusted *R^2^*=-.05) nor for BA_ACC_ indices (*F*_8,16_=1.07, *p*=.429, adjusted *R^2^*=-.023, respectively). (Figure 11A, B).

**Figure 11.**
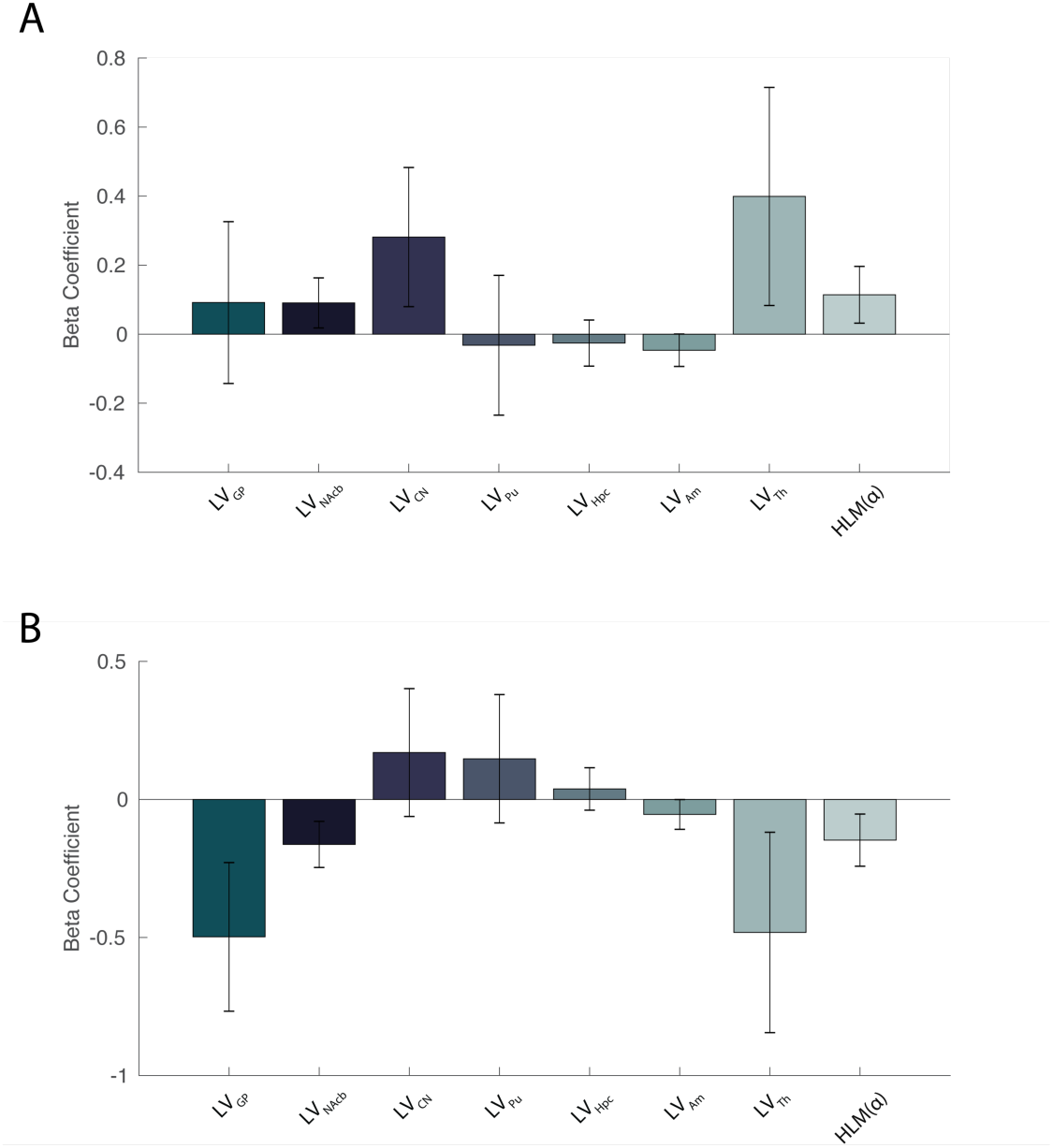
General linear model displaying combined lateralized subcortical volumes and hemispheric lateralized modulation as multiple regressors for the prediction of spatial behavioural bias in RT (A) and Accuracy (B). No significant regression was found which could account for either the lateralized accuracy or RTs (*p*=.429 and *p*=.570, respectively).

## Discussion

The aim of this study was to investigate the involvement of subcortical structures in modulating spatial attention to stimuli associated with contextual salience. We observed that volumetric lateralization of subcortical areas explained individual differences in the ability to modulate interhemispheric alpha power. Specifically, participants exhibiting a right lateralized globus pallidus (GP) also had a better ability to modulate posterior alpha oscillations in the right compared to left hemisphere, and vice versa. The same association held for the relationship between thalamic (Th) hemispheric asymmetry and alpha modulation. Importantly, only the correlation between GP and alpha hemispheric lateralized modulation increased as a function of value-saliency occurrences in the task. To the best of our knowledge, this is the first finding relating individual volumetric differences in BG and thalamus to the modulation of posterior alpha oscillations.

### Subcortical areas and alpha synchronization

Our first finding is in line with a growing body of literature demonstrating a subcortical involvement in high level cognitive functions, such as conscious perception (Slagter et al., 2016), working memory performance (Frank et al., 2001), cognitive control (Reilly et al., 2011; Ceaser and Barch, 2015; Piray et al., 2016) and attentional control (Yantis et al., 2012; Tommasi et al., 2015). We showed that volumetric asymmetry of subcortical areas predicts individual biases in the ability to efficiently allocate attention, as indexed by interhemispheric modulation of alpha power. This is strong support in favour of a subcortical involvement in attentional processing, given the well-established role of alpha oscillations in the allocation of spatial visuospatial attention (Jensen and Mazaheri, 2010). In line with the functional association between BG and cognitive control in the context of reward (Fallon and Cools, 2014; Fallon et al., 2017), we provide novel insights into the involvement of subcortical regions in the modulation of posterior alpha oscillations.

### Pulsed inhibition

A well-recognized function of the BG is to inhibit or promote cortical activity via GABAergic signalling, through the globus pallidus pars interna (GPi), one of its major output structures (Lanciego et al., 2012; Goldberg et al., 2013). The BG might exercise its influence by applying control over activity in the prefrontal cortex or it might directly coordinate posterior regions (as reflected by its relationship to alpha power modulation during reward processing). Our results suggest that individual differences in GP volume lateralization may correspond to interhemispheric variability in GABAergic signalling and thus reflect the subcortical potential to inhibit cortex. This input is likely responsible for producing the mechanisms of ‘*pulsed inhibition*’ in the visual cortex (Jensen and Mazaheri, 2010), reflected by interhemispheric modulation of alpha power, allowing the selective processing of stimuli. Implicitly, we assumed that the volume of the GP indirectly reflects its ability to exert its top-down control over posterior areas, its size possibly representing a determinant for the number of GABAergic neurons involved in the control mechanism.

### GP in relation to attentional selection and cognitive control

Interestingly, our results emphasize the specific contribution of the GP in supporting stimulus-driven allocation of attention in a value-based context. The GPi is considered to mediate the output of the BG (Lanciego et al., 2012; Goldberg et al., 2013), and previous literature has implicated this structure in voluntary movement regulation: its functions have indeed been predominantly investigated in clinical and animal models in association with motor functions and action control (Filion and Tremblay, 1991; Jahfari et al., 2011), describing, for instance, reduction of hypokinetic and rigidity symptoms following pallidotomy in humans (Schuurman et al., 1997; Dostrovsky et al., 2002). Nevertheless, recent results from single unit recordings in humans provided indications that electrophysiological activity in the GPi reflects processing of stimuli associated with different reward contingencies (Howell et al., 2016). This is corroborated by evidence of alterations of cognitive, in addition to motor, abilities, following pallidotomy in Parkinson’s disease (PD) patients (Lombardi et al., 2000); In addition, electrical stimulation of the GPi to treat PD has been reported to be associated with several cognitive impairments, such as subtle declines in attention and concentration, although to a lesser extent when compared to subthalamic stimulation (Combs et al., 2015). This aspect has been further addressed in clinical studies showing a link between PD, associated with abnormal pallidal activity (Dostrovsky et al., 2002; Rosenberg-Katz et al., 2016), and altered reward processing as well as updating (Aarts et al., 2012; Chong et al., 2015). Structural GP abnormalities have also been linked to impaired suppression of distractors in ADHD (Aylward et al., 1996; Qiu et al., 2009) and psychotic symptoms in schizophrenia (Hokama et al., 1995; Spinks et al., 2005; Mamah et al., 2007), which has been related to aberrant salience attribution and reward learning (Early et al., 1987; Okada et al., 2016). As an important output component of the reward circuit (Haber, 2011), the GPi might serve to indirectly influence the cortical information flow by biasing selective processing of value-related stimuli. Our data expands on this notion by suggesting a further pallidal influence on the modulation of visual alpha oscillations.

Importantly, given that the association between LV_GP_ and HLM(α) increases as a function of the saliency, we here postulate a specific role for the GP in value-related shifts of attention. On the other hand, it is still unclear to what extent this modulation is dependent on value-related stimuli rather than covert visual attention: additional studies would be valuable to further disentangle the role of these two features and generalize the findings.

### Right lateralization of the association between GP and alpha modulation

Notably, the association between GP lateralization and interhemispheric alpha power (Figure 6) was largely related to right hemisphere differences in absolute alpha modulation between subjects exhibiting a right, as compared to left, lateralized GP volume. This finding possibly reflects the right hemisphere dominance allegedly characterizing spatial attention processes (Shulman et al., 2010b), corroborated by the right lateralized feature of the ventral attentional network, which has been described as specifically involved in the processing of behaviourally salient stimuli (Corbetta and Shulman, 2002)

### Differential role of GP and Th in relation to posterior alpha modulation

Our results show that GP and Th lateralizations were related to the interhemispheric bias in alpha modulation during selective allocation of attention. However, only GP lateralization was related to the value-saliency pairings in the task. The different contribution from GP and Th in relation to saliency occurrences might likely reflect different roles of the two areas in the top-down control of attentional processing. The GP provides a modulatory signal related to the processing of stimuli that draw attention due to their strong saliency associations. The perceptual competition resulting from attending to a salient target whilst required to suppress an equally salient distractor, might be resolved by a network involving the GP. Increased midbrain activity has indeed been shown to accompany attentional suppression of a highly rewarding distractor carrying a strong perceptual competition with the target (Gong et al., 2017), suggesting that dopaminergic networks might flexibly modulate attentional selection in reward-related contexts.

With regard to thalamic regulation of interhemispheric alpha power, it is important to mention that our interpretation is limited by the current pragmatic difficulty in reliably disentangling different thalamic nuclei’s volume, by means of the automatic segmentation algorithm. By considering the full thalamic volume, one is subject to intrinsic confounds derived from the fact that thalamic nuclei might exert differential modulatory effects on cortical activity. It is not to be excluded that a saliency specific processing might still occur within specific nuclei in the structure.

In our sample, the correlation between thalamic lateralization and attention-related alpha modulation was irrespective of the saliency component in the current task. Despite the considerations above, our interpretation of the findings builds upon previous extensive evidence describing how thalamic activity, particularly arising from its largest nucleus, the pulvinar, modulates the alpha rhythm in extended visual areas (Lopes da Silva et al., 1980; Wilke et al., 2009; Saalmann et al., 2012; Zhou et al., 2016; Green et al., 2017). The pulvinar was first shown to contribute to the generation of the posterior alpha rhythm in dogs (Lopes da Silva et al., 1980) and also to regulate synchronized activity between visual cortical areas to support the allocation of attention in human and nonhuman primates (Petersen et al., 1987; Kastner et al., 2004; Saalmann et al., 2012; Green et al., 2017). Our findings, therefore, add to the growing body of evidence suggesting that thalamo-cortical interactions play a fundamental role in shaping cognitive processing (Saalmann and Kastner, 2011; Leszczyński and Staudigl, 2016; Sherman, 2016; Green et al., 2017; Halassa and Kastner, 2017; Fiebelkorn and Kastner, 2019).

### Pallido-cortical pathways

Through which route does the GP influence visual alpha oscillations? A possibility is that the GP modulates prefrontal activity which in turn engages and affects dorsal attentional networks (Cummings, 1993; Pauls et al., 2014). The dorsal attention network, with the intraparietal sulcus (IPS) and frontal eye-fields (FEF) as its major hubs, has been suggested to mediate top-down allocation of attention. Both the IPS and FEF have been indeed causally implicated in the control over posterior alpha oscillations in relation to attentional shifts (Corbetta and Shulman, 2002; Capotosto et al., 2012b; Ptak, 2012; Vossel et al., 2014; Marshall et al., 2015a). Based on our results, we propose the existence of a network which allows salience driven signals from the BG to influence the prefrontal cortex in biasing the competition among posterior regions. The idea of a BG-cortico loop involved in stimulus driven reorienting of attention has been already introduced (Alexander, 1986; Shulman et al., 2010a) and is consistent with the notion of a ‘salience network’, which integrates behaviourally relevant input in order to bias and guide cognitive control (Seeley et al., 2007; Metzger, 2010; Chen et al., 2015; Peters et al., 2016). Within this framework, the BG, through their main output via the GPi, are thought to influence the connectivity between frontoparietal regions by updating goal-directed behaviour, in order to adapt to changes in the environment (van Schouwenburg et al., 2010b).

The influence of GP on posterior alpha oscillations could further be mediated through indirect projections via the thalamus. The major target of GPi projections is the motor thalamus, including ventrolateral and ventral anterior thalamic nuclei, which innervates motor and premotor cortex (Herrero et al., 2002; Sommer, 2003; Goldberg et al., 2013). However, cortical projections from thalamic nuclei receiving input from the BG might be more diverse and target also prefrontal areas (McFarland and Haber, 2002), which would enable an indirect modulation of frontoparietal networks by the GPi via the thalamus. Additionally, intra-thalamic connectivity (Crabtree et al., 1998; Crabtree and Isaac, 2002) as well as complex interactions between the thalamic reticular nucleus and thalamic nuclei (Guillery et al., 1998; Halassa and Acsády, 2016) may provide multiple alternative pathways to convey influence of the GPi on cortical areas and modulate behaviour (Haber and Calzavara, 2009).

The proposed models provide a theoretical framework in favour of a flexible subcortical modulation of top-down regulation of attentional allocation, which for the GP appears to be specifically involved in tasks involving value-saliency processing. Nevertheless, the aforementioned possible modulatory routes should not be considered as mutually exclusive: a more comprehensive model of attentional control should instead account for multiple cortical and subcortical pathways operating in parallel, which would allow optimization of the organism’s interaction with the environment.

### Data availability

The preprocessed MEG and MRI anonymised datasets that support the findings of this study are available as downloadable online data collection in the Donders Data Repository (https://data.donders.ru.nl), with persistent identifier: <u>11633/di.dccn.DSC_3016045.01_337</u>, upon reasonable request to the corresponding author.

## Acknowledgements

The authors gratefully acknowledge the support of the Netherlands Organisation for Scientific Research (NWO, VICI grants 453-09-002 and 453-14-015 and the James S. McDonnell Foundation (grants 220020328 and 220020448). We also would like to thank Sebastiaan den Boer for his in contribution in the data collection process and experimental design. SJF was supported by the NIHR Biomedical Research Centre at University Hospitals Bristol NHS Foundation Trust and the University of Bristol. The views expressed in this publication are those of the author(s) and not necessarily those of the NHS, the National Institute for Health Research or the Department of Health and Social Care.

## References

Aarts E, Helmich RC, Janssen MJR, Oyen WJG, Bloem BR, Cools R (2012) Aberrant reward processing in Parkinson’s disease is associated with dopamine cell loss. Neuroimage 59:3339–3346.

Alexander G (1986) Parallel Organization of Functionally Segregated Circuits Linking Basal Ganglia and Cortex. Annu Rev Neurosci 9:357–381.

Allen M, Poggiali D, Whitaker K, Marshall TR, Kievit R (2018) Raincloud plots: a multi-platform tool for robust data visualization. PeerJ Prepr 6:e27137v1.

Arcizet F, Krauzlis RJ (2018) Covert spatial selection in primate basal ganglia. PLoS Biol 16:e2005930.

Aylward EH, Reiss AL, Reader MJ, Singer HS, Brown JE, Denckla MB (1996) Basal ganglia volumes in children with attention-deficit hyperactivity disorder. J Child Neurol 11:112–115.

Bastiaansen MCM, Knösche TR (2000) Tangential derivative mapping of axial MEG applied to event-related desynchronization research. Clin Neurophysiol 111:1300–1305.

Bischoff-Grethe A, Ozyurt IB, Busa E, Quinn BT, Fennema-Notestine C, Clark CP, Morris S, Bondi MW, Jernigan TL, Dale AM, Brown GG, Fischl B (2007) A technique for the deidentification of structural brain MR images. Hum Brain Mapp 28:892–903.

Braunlich K, Seger C (2013) The basal ganglia. Wiley Interdiscip Rev Cogn Sci 4:135–148.

Capotosto P, Corbetta M, Romani GL, Babiloni C (2012a) Electrophysiological correlates of stimulus-driven reorienting deficits after interference with right parietal cortex during a spatial attention task: a TMS-EEG study. J Cogn Neurosci 24:10.1162/jocn_a_00287.

Capotosto P, Corbetta M, Romani GL, Babiloni C (2012b) Electrophysiological correlates of stimulus-driven reorienting deficits after interference with right parietal cortex during a spatial attention task: a TMS-EEG study. J Cogn Neurosci 24:10.1162/jocn_a_00287.

Ceaser AE, Barch DM (2015) Striatal Activity is Associated with Deficits of Cognitive Control and Aberrant Salience for Patients with Schizophrenia. Front Hum Neurosci 9:687.

Chelazzi L, Perlato A, Santandrea E, Della Libera C (2013) Rewards teach visual selective attention. Vision Res 85:58–62.

Chen MC, Ferrari L, Sacchet MD, Foland-Ross LC, Qiu MH, Gotlib IH, Fuller PM, Arrigoni E, Lu J (2015) Identification of a direct GABAergic pallidocortical pathway in rodents. Eur J Neurosci 41:748–759.

Chong TTJ, Bonnelle V, Manohar S, Veromann KR, Muhammed K, Tofaris GK, Hu M, Husain M (2015) Dopamine enhances willingness to exert effort for reward in Parkinson’s disease. Cortex 69:40–46.

Combs HL, Folley BS, Berry DTR, Segerstrom SC, Han DY, Anderson-Mooney AJ, Walls BD, van Horne C (2015) Cognition and Depression Following Deep Brain Stimulation of the Subthalamic Nucleus and Globus Pallidus Pars Internus in Parkinson’s Disease: A Meta-Analysis. Neuropsychol Rev 25:439–454.

Corbetta M, Shulman GL (2002) Control of Goal-Directed and Stimulus-Driven Attention in the Brain. Nat Rev Neurosci 3:215–229.

Crabtree JW, Collingridge GL, Isaac JTR (1998) A new intrathalamic pathway linking modality-related nuclei in the dorsal thalamus. Nat Neurosci 1:389–394.

Crabtree JW, Isaac JTR (2002) New intrathalamic pathways allowing modality-related and cross-modality switching in the dorsal thalamus. J Neurosci 22:8754–8761.

Cummings J (1993) Frontal-subcortical circuits and human behavior. Arch Neurol 50:873–880.

Dostrovsky JO, Hutchison WD, Lozano AM (2002) The Globus Pallidus, Deep Brain Stimulation, and Parkinson’s Disease. Neurosci 8:284–290.

Early TS, Reiman EM, Raichle ME, Spitznagel EL (1987) Left globus pallidus abnormality in never-medicated patients with schizophrenia. Proc Natl Acad Sci U S A 84:561–563.

Fallon SJ, Cools R (2014) Reward acts on the pFC to enhance distractor resistance of working memory representations. J Cogn Neurosci 26:2812–2826.

Fallon SJ, Zokaei N, Norbury A, Manohar SG, Husain M (2017) Dopamine Alters the Fidelity of Working Memory Representations according to Attentional Demands. J Cogn Neurosci 29:728–738.

Fiebelkorn IC, Kastner S (2019) The Puzzling Pulvinar. Neuron 101:201–203.

Fiebelkorn IC, Pinsk MA, Kastner S (2019) The mediodorsal pulvinar coordinates the macaque fronto-parietal network during rhythmic spatial attention. Nat Commun 10.

Filion M, Tremblay L (1991) Abnormal spontaneous activity of globus pallidus neurons in monkeys with MPTP-induced parkinsonism. Brain Res 547:140–144.

Frank MJ, Loughry B, O’Reilly RC (2001) Interactions between frontal cortex and basal ganglia in working memory: A computational model. Cogn Affect Behav Neurosci 1:137–160.

Goldberg JH, Farries MA, Fee MS (2013) Basal ganglia output to the thalamus: still a paradox. Trends Neurosci 36:10.1016/j.tins.2013.09.001.

Gong M, Jia K, Li S (2017) Perceptual Competition Promotes Suppression of Reward Salience in Behavioral Selection and Neural Representation. J Neurosci 37:6242–6252.

Green JJ, Boehler CN, Roberts KC, Chen L-C, Krebs RM, Song AW, Woldorff MG (2017) Cortical and Subcortical Coordination of Visual Spatial Attention Revealed by Simultaneous EEG–fMRI Recording. J Neurosci 37:7803–7810.

Guadalupe T et al. (2016) Human subcortical brain asymmetries in 15,847 people worldwide reveal effects of age and sex. Brain Imaging Behav:1–18.

Guillery RW, Feig SL, Lozsádi D a., Lozsadi DA, Lozsádi D a. (1998) Paying attention to the thalamic reticular nucleus. Trends Neurosci 21:28–32.

Haber S (2011) Neuroanatomy of Reward: A View from the Ventral Striatum. In: Neurobiology of Sensation and Reward (Jay A. Gottfried, ed). CRC Press.

Haber SN, Calzavara R (2009) The cortico-basal ganglia integrative network: The role of the thalamus. Brain Res Bull 78:69–74.

Halassa MM, Acsády L (2016) Thalamic Inhibition: Diverse Sources, Diverse Scales. Trends Neurosci 39:680–693.

Halassa MM, Kastner S (2017) Thalamic functions in distributed cognitive control. Nat Neurosci 20:1669–1679.

Halgren M, Devinsky O, Doyle WK, Bastuji H, Rey M, Mak-McCully R, Chauvel P, Ulbert I, Fabó D, Erőss L, Wittner L, Heit G, Eskandar E, Mandell A, Cash SS (2017) The Generation and Propagation of the Human Alpha Rhythm. bioRxiv:202564.

Herrero MT, Barcia C, Navarro JM (2002) Functional anatomy of thalamus and basal ganglia. Child’s Nerv Syst 18:386–404.

Hikosaka O, Bromberg-Martin E, Hong S, Matsumoto M (2008) New insights on the subcortical representation of reward. Curr Opin Neurobiol 18:203–208.

Hikosaka O, Kim HF, Yasuda M, Yamamoto S (2014) Basal Ganglia Circuits for Reward Value– Guided Behavior. Annu Rev Neurosci 37:289–306.

Hokama H, Shenton ME, Nestor PG, Kikinis R, Levitt JJ, Metcalf D, Wible CG, O’Donnella BF, Jolesz FA, McCarley RW (1995) Caudate, putamen, and globus pallidus volume in schizophrenia: A quantitative MRI study. Psychiatry Res Neuroimaging 61:209–229.

Howell NA, Prescott IA, Lozano AM, Hodaie M, Voon V, Hutchison WD (2016) Preliminary evidence for human globus pallidus pars interna neurons signaling reward and sensory stimuli. Neuroscience 328:30–39.

Jahfari S, Waldorp L, van den Wildenberg WPM, Scholte HS, Ridderinkhof KR, Forstmann BU (2011) Effective Connectivity Reveals Important Roles for Both the Hyperdirect (Fronto-Subthalamic) and the Indirect (Fronto-Striatal-Pallidal) Fronto-Basal Ganglia Pathways during Response Inhibition. J Neurosci 31:6891–6899.

Jaramillo J, Mejias JF, Wang XJ (2019) Engagement of Pulvino-cortical Feedforward and Feedback Pathways in Cognitive Computations. Neuron 101:321–336.e9.

Jensen O, Mazaheri A (2010) Shaping functional architecture by oscillatory alpha activity: gating by inhibition. Front Hum Neurosci 4:186.

Kastner S, O’Connor DH, Fukui MM, Fehd HM, Herwig U, Pinsk MA (2004) Functional imaging of the human lateral geniculate nucleus and pulvinar. J Neurophysiol 91:438–448.

Kelly SP (2006) Increases in Alpha Oscillatory Power Reflect an Active Retinotopic Mechanism for Distracter Suppression During Sustained Visuospatial Attention. J Neurophysiol 95:3844–3851.

Lanciego JL, Luquin N, Obeso JA (2012) Functional neuroanatomy of the basal ganglia. Cold Spring Harb Perspect Med 2:1–20.

Lauwereyns J, Takikawa Y, Kawagoe R, Kobayashi S, Koizumi M, Coe B, Sakagami M, Hikosaka O (2002) Feature-based anticipation of cues that predict reward in monkey caudate nucleus. Neuron 33:463–473.

Leszczyński M, Staudigl T (2016) Memory-guided attention in the anterior thalamus. Neurosci Biobehav Rev 66:163–165.

Lombardi WJ, Gross RE, Trepanier LL, Lang AE, Lozano AM, Saint-Cyr JA (2000) Relationship of lesion location to cognitive outcome following microelectrode-guided pallidotomy for Parkinson’s disease: support for the existence of cognitive circuits in the human pallidum. Brain 123:746–758.

Lopes da Silva FH, Vos JE, Mooibroek J, van Rotterdam A (1980) Relative contributions of intracortical and thalamo-cortical processes in the generation of alpha rhythms, revealed by partial coherence analysis. Electroencephalogr Clin Neurophysiol 50:449–456.

Mamah D, Wang L, Barch D, de Erausquin GA, Gado M, Csernansky JG (2007) Structural analysis of the basal ganglia in schizophrenia. Schizophr Res 89:59–71.

Maris E, Oostenveld R (2007) Nonparametric statistical testing of EEG- and MEG-data. J Neurosci Methods 164:177–190.

Marshall TR, Bergmann TO, Jensen O (2015a) Frontoparietal Structural Connectivity Mediates the Top-Down Control of Neuronal Synchronization Associated with Selective Attention. PLoS Biol 13:1–17.

Marshall TR, den Boer S, Cools R, Jensen O, Fallon SJ, Zumer JM (2017) Occipital Alpha and Gamma Oscillations Support Complementary Mechanisms for Processing Stimulus Value Associations. J Cogn Neurosci:1–11.

Marshall TR, O’Shea J, Jensen O, Bergmann TO (2015b) Frontal Eye Fields Control Attentional Modulation of Alpha and Gamma Oscillations in Contralateral Occipitoparietal Cortex. J Neurosci 35:1638–1647.

McFarland NR, Haber SN (2002) Thalamic relay nuclei of the basal ganglia form both reciprocal and nonreciprocal cortical connections, linking multiple frontal cortical areas. J Neurosci 22:8117–8132.

Metzger CD (2010) High field fMRI reveals thalamocortical integration of segregated cognitive and emotional processing in mediodorsal and intralaminar thalamic nuclei. Front Neuroanat 4:1–17.

Nobre AC (Kia), Kastner S (2014) The Oxford handbook of attention. Oxford: Oxford University Press.

Okada N et al. (2016) Abnormal asymmetries in subcortical brain volume in schizophrenia. Mol Psychiatry 21:1–7.

Oostenveld R, Fries P, Maris E, Schoffelen JM (2011) FieldTrip: Open source software for advanced analysis of MEG, EEG, and invasive electrophysiological data. Comput Intell Neurosci 2011.

Paton JJ, Belova MA, Morrison SE, Salzman CD (2006) The primate amygdala represents the positive and negative value of visual stimuli during learning. Nature 439:865–870.

Pauls DL, Abramovitch A, Rauch SL, Geller DA (2014) Obsessive–compulsive disorder: an integrative genetic and neurobiological perspective. Nat Rev Neurosci 15:410–424.

Peters SK, Dunlop K, Downar J (2016) Cortico-Striatal-Thalamic Loop Circuits of the Salience Network: A Central Pathway in Psychiatric Disease and Treatment. Front Syst Neurosci 10:1–23.

Petersen SE, Robinson DL, Morris JD (1987) Contributions of the pulvinar to visual spatial attention. Neuropsychologia 25:97–105.

Piray P, Toni I, Cools R (2016) Human Choice Strategy Varies with Anatomical Projections from Ventromedial Prefrontal Cortex to Medial Striatum. J Neurosci 36:2857–2867.

Ptak R (2012) The Frontoparietal Attention Network of the Human Brain. Neurosci 18:502–515.

Qiu A, Crocetti D, Adler M, Mahone EM, Denckla MB, Miller MI, Mostofsky SH (2009) Basal ganglia volume and shape in children with attention deficit hyperactivity disorder. Am J Psychiatry 166:74–82.

Reilly RCO, Herd SA, Pauli WM (2011) Computational Models of Cognitive Control. Psychology 20:257–261.

Rosenberg-Katz K, Herman T, Jacob Y, Kliper E, Giladi N, Hausdorff JM (2016) Subcortical Volumes Differ in Parkinson’s Disease Motor Subtypes: New Insights into the Pathophysiology of Disparate Symptoms. Front Hum Neurosci 10:1–9.

Saalmann YB, Kastner S (2011) Cognitive and Perceptual Functions of the Visual Thalamus. Neuron 71:209–223.

Saalmann YB, Pinsk MA, Wang L, Li X, Kastner S (2012) The Pulvinar Regulates Information Transmission Between Cortical Areas Based on Attention Demands. Science (80-) 337:753–756.

Schechtman E, Noblejas MI, Mizrahi AD, Dauber O, Bergman H (2016) Pallidal spiking activity reflects learning dynamics and predicts performance. Proc Natl Acad Sci 113:E6281–E6289.

Schultz W, Tremblay L, Hollerman JR, Schultz (2000) Reward processing in primate orbitofrontal cortex and basal ganglia. Cereb Cortex 10:272–284.

Schuurman PR, de Bie RMA, Speelman JD, Bosch DA (1997) Posteroventral Pallidotomy in Movement Disorders. In: Advances in Stereotactic and Functional Neurosurgery 12: Proceedings of the 12th Meeting of the European Society for Stereotactic and Functional Neurosurgery, Milan 1996 (Ostertag CB, Thomas DGT, Bosch A, Linderoth B, Broggi G, eds), pp 14–17. Vienna: Springer Vienna.

Seeley WW, Menon V, Schatzberg AF, Keller J, Glover GH, Kenna H, Reiss AL, Greicius MD (2007) Dissociable Intrinsic Connectivity Networks for Salience Processing and Executive Control. J Neurosci 27:2349–2356.

Sherman SM (2016) Thalamus plays a central role in ongoing cortical functioning. Nat Neurosci 19:533–541.

Shipp S (2004) The brain circuitry of attention. Trends Cogn Sci 8:223–230.

Shulman GL, Astafiev S V, Franke D, Pope DLW, Abraham Z, Mcavoy MP, Corbetta M (2010a) Interaction of stimulus-driven reorienting and expectation in ventral and dorsal fronto-parietal and basal ganglia-cortical networks. 29:4392–4407.

Shulman GL, Pope DLW, Astafiev S V., McAvoy MP, Snyder AZ, Corbetta M (2010b) Right Hemisphere Dominance during Spatial Selective Attention and Target Detection Occurs Outside the Dorsal Frontoparietal Network. J Neurosci 30:3640–3651.

Slagter HA, Mazaheri A, Reteig LC, Smolders R, Figee M, Mantione M, Schuurman PR, Denys D (2016) Contributions of the ventral striatum to conscious perception: An intracranial EEG study of the attentional blink. J Neurosci 37:1081–1089.

Sommer MA (2003) The role of the thalamus in motor control. Curr Opin Neurobiol 13:663–670.

Spinks R, Nopoulos P, Ward J, Fuller R, Magnotta VA, Andreasen NC (2005) Globus pallidus volume is related to symptom severity in neuroleptic naive patients with schizophrenia. Schizophr Res 73:229–233.

Stolk A, Todorovic A, Schoffelen JM, Oostenveld R (2013) Online and offline tools for head movement compensation in MEG. Neuroimage 68:39–48.

Thut G (2006) -Band Electroencephalographic Activity over Occipital Cortex Indexes Visuospatial Attention Bias and Predicts Visual Target Detection. J Neurosci 26:9494–9502.

Tomer R, Goldstein RZ, Wang G-J, Wong C, Volkow ND (2008) Incentive motivation is associated with striatal dopamine asymmetry. Biol Psychol 77:98–101.

Tomer R, Slagter HA, Christian BT, Fox AS, King CR, Murali D, Davidson RJ (2013) Dopamine Asymmetries Predict Orienting Bias in Healthy Individuals. Cereb Cortex 23:2899–2904.

Tommasi G, Fiorio M, Yelnik J, Krack P, Sala F, Schmitt E, Fraix V, Bertolasi L, Le Bas J-F, Ricciardi GK, Fiaschi A, Theeuwes J, Pollak P, Chelazzi L (2014) Disentangling the Role of Cortico-Basal Ganglia Loops in Top–Down and Bottom–Up Visual Attention: An Investigation of Attention Deficits in Parkinson Disease. J Cogn Neurosci 27:1215–1237.

Tommasi G, Fiorio M, Yelnik J, Krack P, Sala F, Schmitt E, Fraix V, Bertolasi L, Le Bas J-F, Ricciardi GK, Fiaschi A, Theeuwes J, Pollak P, Chelazzi L (2015) Disentangling the Role of Cortico-Basal Ganglia Loops in Top–Down and Bottom–Up Visual Attention: An Investigation of Attention Deficits in Parkinson Disease. J Cogn Neurosci 27:1215–1237.

Tremblay L, Hollerman JR, Schultz W (1998) Modifications of reward expectation-related neuronal activity during learning in primate striatum. J Neurophysiol 80:964–977.

van Schouwenburg M, Aarts E, Cools R (2010a) Dopaminergic modulation of cognitive control: distinct roles for the prefrontal cortex and the basal ganglia. Curr Pharm Des 16:2026–2032.

van Schouwenburg MR, den Ouden HEM, Cools R (2010b) The Human Basal Ganglia Modulate Frontal-Posterior Connectivity during Attention Shifting. J Neurosci 30:9910–9918.

Van Schouwenburg MR, Den Ouden HEM, Cools R (2015) Selective attentional enhancement and inhibition of fronto-posterior connectivity by the basal ganglia during attention switching. Cereb Cortex 25:1527–1534.

Vossel S, Geng JJ, Fink GR (2014) Dorsal and Ventral Attention Systems. Neurosci 20:150–159.

Wilcox RR (2016a) Comparing dependent robust correlations. Br J Math Stat Psychol 69:215–224.

Wilcox RR (2016b) Introduction to Robust Estimation and Hypothesis Testing. Academic Press.

Wilcox RR (2016c) Comparing dependent robust correlations. Br J Math Stat Psychol 69:215–224.

Wilke M, Mueller K-M, Leopold DA (2009) Neural activity in the visual thalamus reflects perceptual suppression. Proc Natl Acad Sci 106:9465–9470.

Womer FY, Wang L, Alpert KI, Smith MJ, Csernansky JG, Barch DM, Mamah D (2014) Basal ganglia and thalamic morphology in schizophrenia and bipolar disorder. Psychiatry Res - Neuroimaging 223:75–83.

Worden MS, Foxe JJ, Wang N, Simpson G V (2000) Anticipatory biasing of visuospatial attention indexed by retinotopically specific alpha-band electroencephalography increases over occipital cortex. J Neurosci 20:RC63.

Yantis S, Anderson BA, Wampler EK, Laurent PA (2012) Reward and attentional control in visual search. Nebr Symp Motiv 59:91–116.

Zheng J, Anderson KL, Leal SL, Shestyuk A, Gulsen G, Mnatsakanyan L, Vadera S, Hsu FPK, Yassa MA, Knight RT, Lin JJ (2017) Amygdala-hippocampal dynamics during salient information processing. Nat Commun 8:1–11.

Zhou H, Schafer RJ, Desimone R (2016) Pulvinar-Cortex Interactions in Vision and Attention. Neuron 89:209–220.

